# Using real-time fluorescence and deformability cytometry and deep learning to transfer molecular specificity to label-free sorting

**DOI:** 10.1101/862227

**Authors:** Ahmad Ahsan Nawaz, Marta Urbanska, Maik Herbig, Martin Nötzel, Martin Kräter, Philipp Rosendahl, Christoph Herold, Nicole Toepfner, Marketa Kubankova, Ruchi Goswami, Shada Abuhattum, Felix Reichel, Paul Müller, Anna Taubenberger, Salvatore Girardo, Angela Jacobi, Jochen Guck

## Abstract

The identification and separation of specific cells from heterogeneous populations is an essential prerequisite for further analysis or use. Conventional passive and active separation approaches rely on fluorescent or magnetic tags introduced to the cells of interest through molecular markers. Such labeling is time- and cost-intensive, can alter cellular properties, and might be incompatible with subsequent use, for example, in transplantation. Alternative label-free approaches utilizing morphological or mechanical features are attractive, but lack molecular specificity. Here we combine image-based real-time fluorescence and deformability cytometry (RT-FDC) with downstream cell sorting using standing surface acoustic waves (SSAW). We demonstrate basic sorting capabilities of the device by separating cell mimics and blood cell types based on fluorescence as well as deformability and other image parameters. The identification of blood sub-populations is enhanced by flow alignment and deformation of cells in the microfluidic channel constriction. In addition, the classification of blood cells using established fluorescence-based markers provides hundreds of thousands of labeled cell images used to train a deep neural network. The trained algorithm, with latency optimized to below 1 ms, is then used to identify and sort unlabeled blood cells at rates of 100 cells/sec. This approach transfers molecular specificity into label-free sorting and opens up new possibilities for basic biological research and clinical therapeutic applications.

## Introduction

The identification and sorting of cells of interest from heterogeneous populations is a necessary first step in many biological, biotechnological, or medical procedures^1–3^. The sorted cells can subsequently be subjected to refined analysis and inquiry into their proteomic, transcriptomic, or genetic identity and function^4,5^. Alternatively, they can be used for culture and serve the establishment of specific drugs^4,6,7^, or for transplantation into patients in regenerative medicine applications^8–12^.

Conventional sorting techniques used for this purpose can be divided into bulk sorters, such as magnetic-activated cell sorting (MACS)^13,14^, filtration-based approaches^15–17^, deterministic lateral displacement^18,19^ or optical lattices^20^, and single-cell sorters, such as fluorescence-activated cell sorter (FACS)^21–23^, as well as light scattering- and imagebased flow cytometers^24–26^. Bulk sorters separate cells passively according to a small number of parameters hard-wired into the experimental setup, can operate in a massively parallel way and have, thus, superior throughput. However, with their use it is not possible to separate particular cells based on a variable choice of orthogonal, individual features (e.g. fluorescent, small, and deformable) in a single sorting run. Single-cell sorters assess cells serially, and are thus slower, but the sorting decision can be made for each cell separately, using features flexibly chosen from a large set. Both bulk and single-cell sorters most often rely on cell identification, and in the case of MACS also cell separation, based on molecular labels, which confer great specificity. Such reliance on molecular labels, however, can alter cells and their function, comes with additional cost and preparation time, renders the experimenter blind to cells without known molecular marker or to subpopulations not yet known, and might be incompatible with subsequent use, for example in transplantation.

Label-free approaches to identify cells promise to address the aforementioned issues. A flagship example is the usage of forward and side scatter signals in flow cytometry to report on the size and internal structure of cells, albeit only in an indirect way^24,27^. Other label-free approaches interrogate diverse chemical or physical properties of cells. Raman scattering, for example, provides multiplexed information on the presence of chemical species in cells^28,29^, quantitative phase imaging yields internal mass density distributions^30–33^, and Brillouin scattering^34,35^ and deformability cytometry^36^ are used for mechanical phenotyping of cells. The latter is particularly relevant to feel for functional changes of cells involving cytoskeletal adaptations^37,38^ or to identify disease states in blood^39^ and can analyze cells in the context of cell circulation and migration ability^40^. One recent implementation of deformability cytometry, which is based on imaging cells flowed through a narrow constriction channel, analyzes the ensuing cell deformation continuously in real-time at rates of 100–1,000 cells/sec^41^, and is thus well-suited for downstream sorting. The image analysis aspect is similar to commercial imaging flow cytometry (IFC), where 2D spatial bright-field and fluorescence information is obtained in high-throughput^42^. While using fluorescence information to identify cells has the draw-backs discussed above, its correlation with bright-field images and cell morphology in IFC is intriguing^25,43^. Although IFC is commercially established^42^ and has opened a steadily growing field of applications^25,43,44^, its extension to sorting is technically challenging as it necessitates real-time image processing and advanced hardware solutions, and has, thus, only been demonstrated in few highly specialized laboratories^25,26^. An important aspect of IFC is the provision of huge numbers of cell images, which predestines it to be combined with artificial intelligence analysis^25,43^.

Here we introduce a robust sorting platform, combining image-based morphological cell analysis with mechanical characterization and subsequent active sorting by feeding real-time fluorescence and deformability cytometry (RT-FDC)^45^ information to a down-stream SSAW-based cell sorter^23^. We demonstrate its ability to identify and sort cell mimics, as well as various blood cells, based on the parameters available from image analysis (size, deformation, brightness), also in combination with fluorescence, at the rates of 100 cells/sec. We further show that the cell passage through a constriction simplifies automated image analysis by aligning cells along their major symmetry axis and enhances the separation of subpopulations by inducing cell deformation. Importantly, since RT-FDC also provides three-channel fluorescence information for each cell in real-time, we can generate training sets of hundreds of thousands of images labeled based on molecular markers in order to train a deep neural network (DNN) for label-free cell identification based on bright-field images alone. We demonstrate this possibility by first training a DNN using a set of cell images classified based on the expression of CD66 and CD14 surface markers, and then using this classifier to sort 50,000 non-stained neutrophils based on the bare cell images. Such identification and sorting, optimized for latency between cell detection and sorting trigger below 1 ms, resulted in 89.7% purity and 10-fold enrichment of neutrophils. This application is of particular relevance since neutrophils are notorious for becoming activated by molecular interventions. Taken together, sorting RT-FDC (soRT-FDC) makes it possible to not only combine the specificity of molecular labeling with the additional information of morphological and mechanical phenotypes, but also to transfer molecular specificity into label-free sorting using artificial intelligence, thereby avoiding the disadvantages of molecular labeling. This new approach of specific and non-invasive cell sorting paves the way to many applications in basic biological research, as well as in diagnostics and regenerative medicine.

## Results

### Combining SSAW-enabled sorting with RT-FDC

RT-FDC is a microfluidic platform that combines high-throughput bright-field imaging of cells with flow cytometry-like acquisition of 1D fluorescence signals in 3 spectral channels^45^. Since the imaging takes place in a narrow channel, it enables extraction of features related not only to cell morphology, but also to cell mechanics. Utility of RT-FDC for basic research as well as for prospective medical use, for example in blood analysis^39^, has been well demonstrated, however, to fully harness the potential of this method, the capability to actively sort cells based on recorded features was missing.

To address this need, and enable sorting based on combinations of cell morphology, mechanics, and fluorescence, we have implemented an SSAW-based sorting capability^23^ to the RT-FDC setup^45^. To this end, a standard RT-FDC chip was modified by adding the sorting region after the analysis channel as depicted in **Figure 1a**. Downstream of the measurement region of interest (red dashed rectangle) the analysis channel widens from 20 or 30 μm to 50 μm to allow for SSAW-induced cell deflection, before it bifurcates into default and target outlets. The channel diameter of 50 μm ensures that, upon excitation of interdigital transducers (IDT) at their designed resonance frequency of ca. 54 MHz (λ = 70 μm), only one pressure node is present within the channel width. The chip consists of a lithium niobate substrate, a PDMS slab with the imprinted channel structures bonded on top, and two IDTs with electrical contacts at each channel side (**Figure 1b**). The phase shift between the two IDT actuators is tuned so that the pressure node is located off-center (**Figure 1b**), which causes a deflection of cells of interest into the target outlet. Due to a slight asymmetry at the bifurcation point (5 μm shift from the channel center) all cells are collected in the default outlet when SSAW is switched off. With SSAW on, the cells are pushed into the pressure node and guided to the target outlet (**Figure 1c**, **Supplementary Figure 1**). For sufficient deflection, cells need to be exposed to the sound waves for about two milliseconds, which is ensured by the applied flow rate (0.04 μl/s or 0.08 μl/s, depending on the size of the analysis channel) and the chosen channel geometry in the sorting region (50 μm width and 200 μm length, height corresponds to the size of the analysis channel). The trigger for SSAW actuation is based on the real-time assessment of high-speed microscopy images or fluorescence signals from one of three available fluorescence channels (FL-1, FL-2, and FL-3; for spectral specifications, and full list of sorting parameters assessed, see **Supplementary Table 1**) in the measurement region of interest (**Figure 1a**). The processing pipeline was implemented in a custom C++ program running on a standard PC. The latency between cell detection and sorting trigger was optimized to be held below one millisecond, which is suitable for reliably analyzing up to 100 cells/s at typical cell concentrations during RT-FDC experiments.

**Figure 1.**
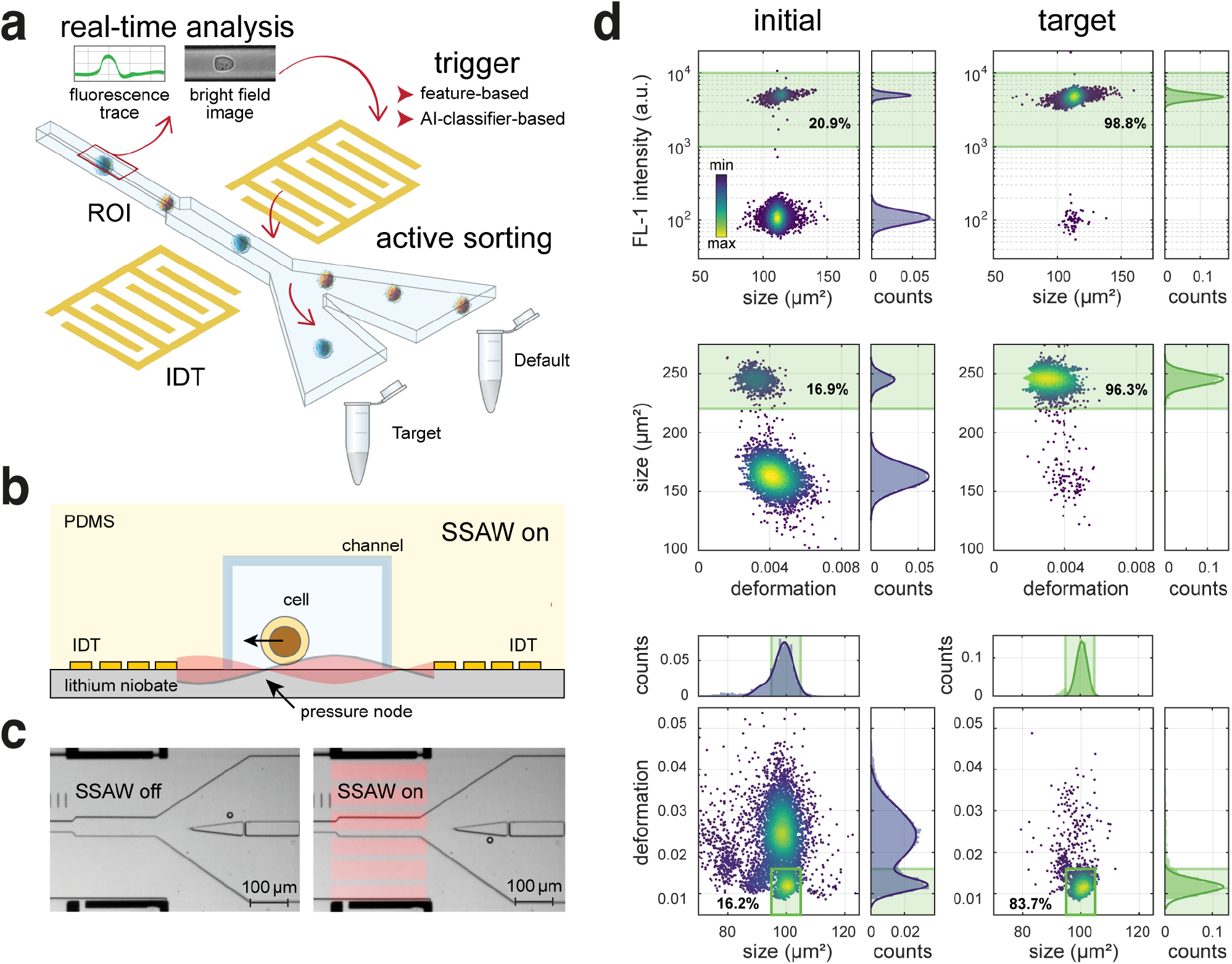
Sorting real-time fluorescence and deformability cytometry (soRT-FDC) is a platform that enables active sorting based on fluorescence and image-derived parameters. (**a**) Operation principle of a soRT-FDC device. Flowed-in cells are deformed in a constriction of the microfluidic channel and analyzed in the region of interest (ROI, outlined in red) in real-time. The fluorescence data as well as bright-field images collected for each cell can be used for making a feature- or artificial intelligence (AI)-classifier-based decision to trigger actuation of a standing surface acoustic wave (SSAW) by interdigital transducers (IDTs). The SSAW deflects the trajectories of selected cells to the target outlet. (**b**) A side view of the device cross-section in the sorting region. In the presence of SSAW, cells are pushed towards the pressure node (indicated with an arrow). (**c**) Micrographs of sorting region of the device taken when SSAW is off (left image, cells flow towards the default outlet), and when SSAW is on (right image, cells are deflected into the target channel). (**d**) Analysis of feature-based sorting of fluorescent and non-fluorescent bead mixture (upper panel), a mixture of beads of two different sizes (middle panel), and a mixture of beads with different stiffnesses (lower panel). On the left, the measurements of initial samples are shown, and on the right the re-analysis of samples collected in the target outlet. The color map in scatter plots represents event density. The histograms of features presented in the scatter plots are shown on top and on the right of the corresponding scatter plots, the histograms were fit with a superposition of Gaussian functions (solid lines). The gates used for sorting are outlined in green. Percentages on scatter plots indicate the fraction of beads in the sorting gate.

### Active sorting using fluorescence and image-derived parameters

To validate the efficacy of SSAW-based sorting in our setup, heterogeneous mixtures of polymer beads — serving as cell mimics — were separated based on features such as fluorescence, size, and deformation.

First, to demonstrate FACS-like capabilities of soRT-FDC to perform fluorescence-activated sorting, we mixed polyacrylamide microgel beads labeled with AlexaFluor488 and unlabeled ones in a 1:5 ratio (beads were produced in house^46^, see **Methods** for details) and sorted for fluorescence intensity (**Figure 1d**, upper panel). The percentage of fluorescence-positive beads contained in the sorting gate (FL-1 intensity between 1,000 and 10,000 a.u.) increased from 20.9% in the initial bead mixture to 98.8% in the sample collected into the target outlet, providing for 4.7-fold enrichment of the labeled beads.

Next, we set out to validate the efficiency of sorting for image-derived parameters such as the object size and deformation. For size-based sorting, we employed a 1:6 mixture of commercially-available monodisperse polymer beads of two different sizes (with a diameter of 13.79 ± 0.59 μm and 17.23 ± 0.24 μm as specified by the supplier; **Figure 1d**, middle panel). The sorting gate was set to 220 – 300 μm^2^ of bead size expressed as crosssectional area (corresponding to a diameter of 16.74 – 19.54 μm). We observed an enrichment of 5.7-fold for the beads of selected size, with 16.9% of total events contained within the sorting gate in the initial sample and 96.3% in the target sample.

It is important to note that precise determination of cell size is relevant for the identification of cell types in heterogeneous populations such as blood^39^ or for investigating basic scientific questions related to e.g. cell size regulation^47^. While the forward scatter (FSC) intensity in flow cytometry typically correlates with cell size, it does not provide a quantitative measure of it and is further influenced by other parameters such as the object’s refractive index^24,27^. In fact, the relative differences in the object sizes captured by a flow cytometer can, in certain cases, be misleading. To illustrate this issue, we have measured a mixture of monodisperse beads of four different diameters made of a different material each using both soRT-FDC and a high-performance flow cytometer (BD LSR II, BD Biosciences) (**Supplementary Figure 2**). With soRT-FDC, we were able to capture relative differences in bead sizes and determine the bead diameter with an overestimation of 2 – 8%, while reproducing the standard deviation of the diameters reported by the supplier. Strikingly, we observed that FSC measurements performed with flow cytometer did not faithfully reflect the differences in bead sizes.

**Figure 2.**
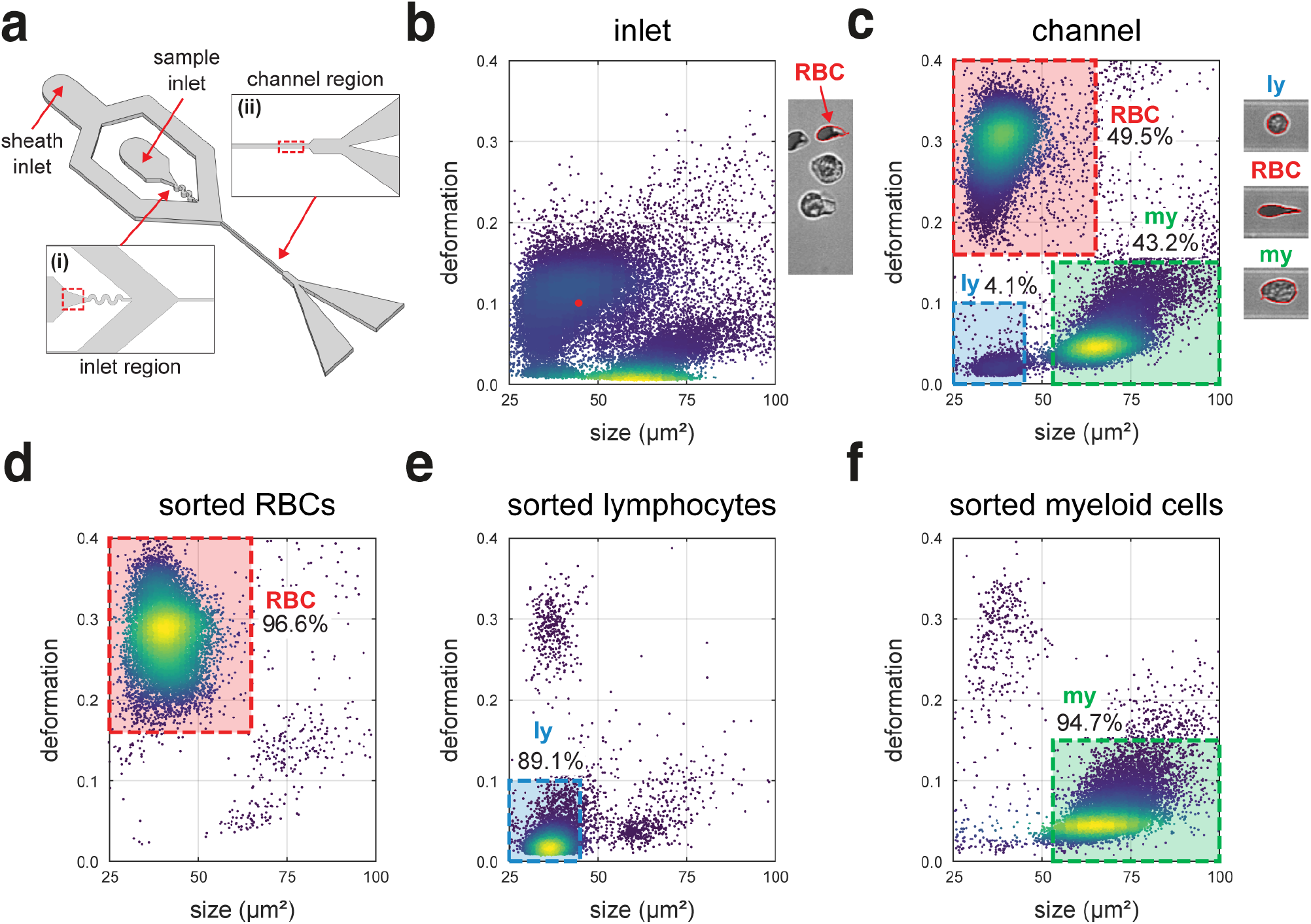
Deformation-assisted sorting for blood cell types. (**a**) Isometric view of the sorting device (not to scale). Inset (i) indicates the sample inlet region where cells are undeformed and randomly oriented. Inset (ii) shows the analysis channel region where cells are deformed and aligned along major channel axis. (**b**) Deformation-size scatter plot of RBC-depleted blood (dextran sedimentation) measured in the inlet region as indicated in (**a**). The red data point in the scatter as well as the arrow on the side indicate a tracked (red periphery around cell) randomly oriented RBC (remaining despite depletion). (**c**) Deformation-size scatter plot of RBC-depleted blood measured in the channel region as shown in (**a**). The indicated cell populations include: red blood cell (RBC, 49.6%), lymphocytes (ly, 4.3%) and myeloid cells (my, 43.1%). The typical images of cells from each population are shown on the right. (**d**) Post analysis of sorted RBCs with 96.6% purity. (**e**) Post analysis of sorted lymphocytes with 89.1% purity. (**f**) Post analysis of sorting for myeloid cell population with 94.7% purity.

Another image-derived parameter that is of high interest for sorting, as it gives access to mechanical properties of cells^48,49^, is deformation. To demonstrate sorting for deformation, we utilized a mixture of two polyacrylamide microgel bead populations of different stiffness^46^. We sorted for the beads with low deformation values of 0.000 – 0.016 contained within the size range of 95 – 105 μm^2^ (**Figure 1d**, lower panel). The percentage of beads in the gate increased from 16.2% in the initial mixture (prepared in a 1:6 ratio), to 83.7% in the target sample, amounting to 5.2-fold enrichment. This experiment illustrates not only sorting for deformation, but also the capability of soRT-FDC to select objects using combinations of features of interest. Further examples of sorting using two orthogonal features, such as florescence intensity and deformation (**Supplementary Figure 3a**), or size and average brightness (**Supplementary Figure 3b**), are presented in **Supplementary Information**.

Sorting of objects based on size and deformation was also demonstrated for living cells. Specifically, we were able to enrich human promyelocytic leukemia cells (HL60/S4) in their mixture with Kc167 drosophila cells from 22.1% to 88.2% through size-based sorting (**Supplementary Figure 4**, **Supplementary Video 1**). Deformation-based sorting allowed for enrichment of red blood cells (RBCs) with high or low deformation values (**Supplementary Figure 5**). These experiments confirm that cells, which are more difficult to translocate using SSAW due to their lower acoustic contrast factor compared to polymer beads^50^, can also be successfully sorted in our setup.

Finally, since it is important for the subsequent use to validate if cells are in good condition after the sorting procedure, we analyzed the viability, proliferation, and mechanical properties of cells after exposure to SSAW. The cells exposed to SSAW in soRT-FDC were able to proliferate at rates similar to that of control cells (**Supplementary Figure 6a**,**b**), their viability in culture remained at levels exceeding 90% (**Supplementary Figure 6c**), and their mechanical properties were not affected as shown by atomic force microscopy indentation measurements (**Supplementary Figure 7**).

### Deformation-assisted sorting for major blood cell types

The efficient separation of cells based on their ability to deform using soRT-FDC opens up interesting experimental avenues in and of itself. Additionally, we have observed that the flow-induced deformation of cells, together with their alignment along the major channel axis, enhances the separation of subpopulations in heterogeneous mixtures of cells, which would be obscured otherwise.

A prominent example of the role of deformation in subpopulation analysis is the identification of various blood cell types. Measurements of RBC-depleted blood in the inlet region of the chip (**Figure 2a**, inset i), where the channel is broad and cells are undeformed and randomly oriented, results in overlaps between the populations of RBCs, lymphocytes, and myeloid cells in a deformation-size scatter plot (**Figure 2b**). In this case, well-defined gates for sorting individual cell subtypes are difficult to define. Acquiring the same measurement in the channel region (**Figure 2a**, inset ii), where cells are deformed and aligned with the channel major axis, results in clear separation of blood cell subtypes (**Figure 2c**), as previously established^39,41^. In particular, the highly-deformed RBC population moves further away from the two remaining cell subtypes on the deformation axis. The now distinct populations can be easily gated and sorted for. Upon sorting for gates specified by deformation and size, we were able to achieve a purity of 96.6% for RBCs (**Figure 2d**), 89.1% for lymphocytes (**Figure 2e**), and 94.7% for myeloid cells (**Figure 2f**), with an enrichment factor of 1.95, 21.7 and 2.0, respectively (see **Supplementary Table 2** for more details).

### Label-free neutrophil sorting based on brightness

Both myeloid and lymphoid cells presented above can be further divided into several distinct subtypes. As demonstrated previously^39^, by adding average brightness of pixels contained within the cell contour as an additional separation parameter, we can further distinguish myeloid cells into neutrophils, monocytes, basophils, and eosinophils (**Figure 3a**). Since brightness is a parameter calculated in real-time by our analysis software, it can be utilized for sorting of the specified myeloid blood subtypes. To demonstrate this possibility, we used a combination of size (50 – 100 μm^2^) and brightness (30 – 40 a.u.) gates to perform label-free sorting of neutrophils from RBC-depleted blood (**Supplementary Video 2**). The percentage of events in the sorting gate increased from 19.4% in the initial sample to 81.5% in the target sample (**Figure 3a**), amounting to 4.2-fold enrichment. To validate the identity of sorted cells, we stained both the initial and the target samples after sorting with a mixture of anti-CD66 and anti-CD14 antibodies and measured them with RT-FDC. We could confirm that while in the initial sample the CD66^+^/CD14^−^ cells corresponding to neutrophils^51^ constituted 24.7% of the population, in the target sample their content increased to 84.7% (**Figure 3b**), corresponding roughly to the percentages in the size-brightness sorting gate (**Figure 3a**). 74.4% of all CD66^+^/CD14^−^ events in the initial sample fell into the size-brightness gate specified for vital, resting, single neutrophils^52^, as displayed in the color-coded scatter plots in **Figure 3c**. The remaining CD66^+^/CD14^−^ events appear to belong to other blood cell type populations based on their physical appearance (**Figure 3c**), implicating occurrence of false positive neutrophil classification when using molecular labels. This result shows that the use of physical parameters, such as size and brightness, for cell identification can reduce the false positive rates inherent to fluorescent staining.

**Figure 3.**
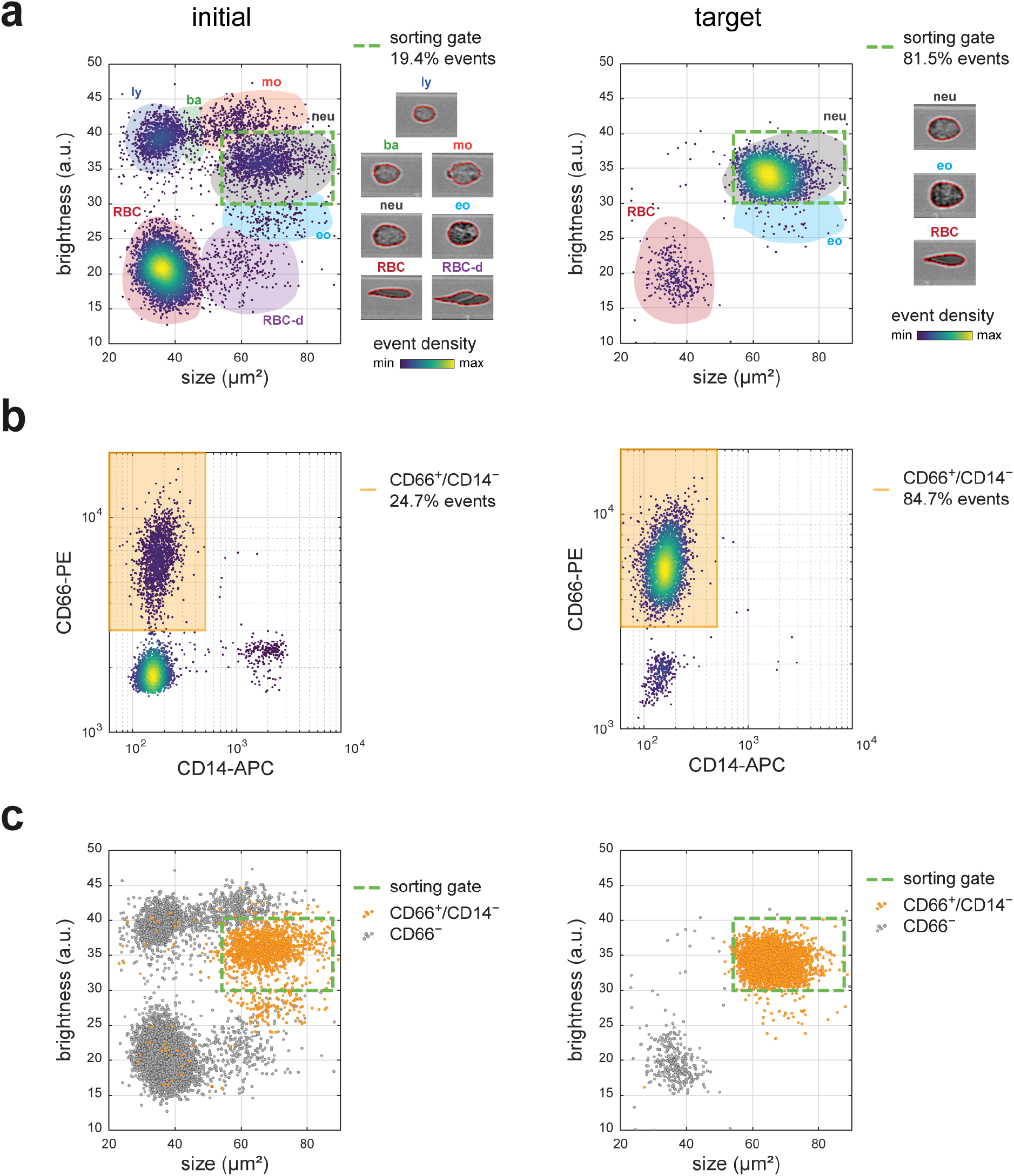
Brightness-based sorting of neutrophils from RBC-depleted blood. (**a**) Brightness-size scatter plots of RBC-depleted blood (dextran sedimentation) for initial (left-hand side) and target (right-hand side) samples. Color-coded patches delineate subpopulations of different cell types. Sorting gate is indicated with dashed green line. Typical cell images of respective subpopulations are displayed next to the scatter plots. The subpopulations include: lymphocytes (ly), basophils (ba), monocytes (mo), neutrophils (neu), eosinophils (eo), red blood cells (RBC) and red blood cell doublets (RBC-d). (**b**) CD66 and CD14 surface marker expression for initial (left) and target (right) samples measured with RT-FDC. (**c**) Brightness-cell size scatter plots as in (**a**) with CD66^+^/CD14^−^ cells (putative neutrophils) indicated in orange.

### DNN-assisted identification and sorting of neutrophils

The image-derived parameters exploited for sorting in the aforementioned examples (such as size, deformation, and brightness) are far from being exhaustive. In fact, there is a plethora of image-based features and their combinations that can be further explored for potential use in cell classification. Examples include Haralick texture features, scale-invariant feature transforms, local binary patterns, or threshold adjacency statistics^53–56^. However, development of such parameters — also known as feature engineering — as well as their selection and classifier training, albeit potentially powerful^57,58^, is typically too computationally expensive to allow for real-time evaluation and applicability for sorting. Another approach is to take advantage of information contained within all pixels of the image, and to use a labeled dataset to train a DNN to classify cells of interest in a featureless way (**Figure 4a**). Supervised machine learning eases the engineering of a predictor for a specific classification task and DNNs have already been shown to outperform traditional handcrafted features and classifiers for different applications^59,60^. DNN training generally requires a sufficiently large training dataset for its successful application. Since RT-FDC can acquire thousands of cell images within a span of seconds and simultaneously evaluate fluorescent antibodies against specific surface markers to provide labels of cell identity for training, our method is primed for the implementation of featureless DNN-based classifiers for label-free cell identification and sorting.

**Figure 4.**
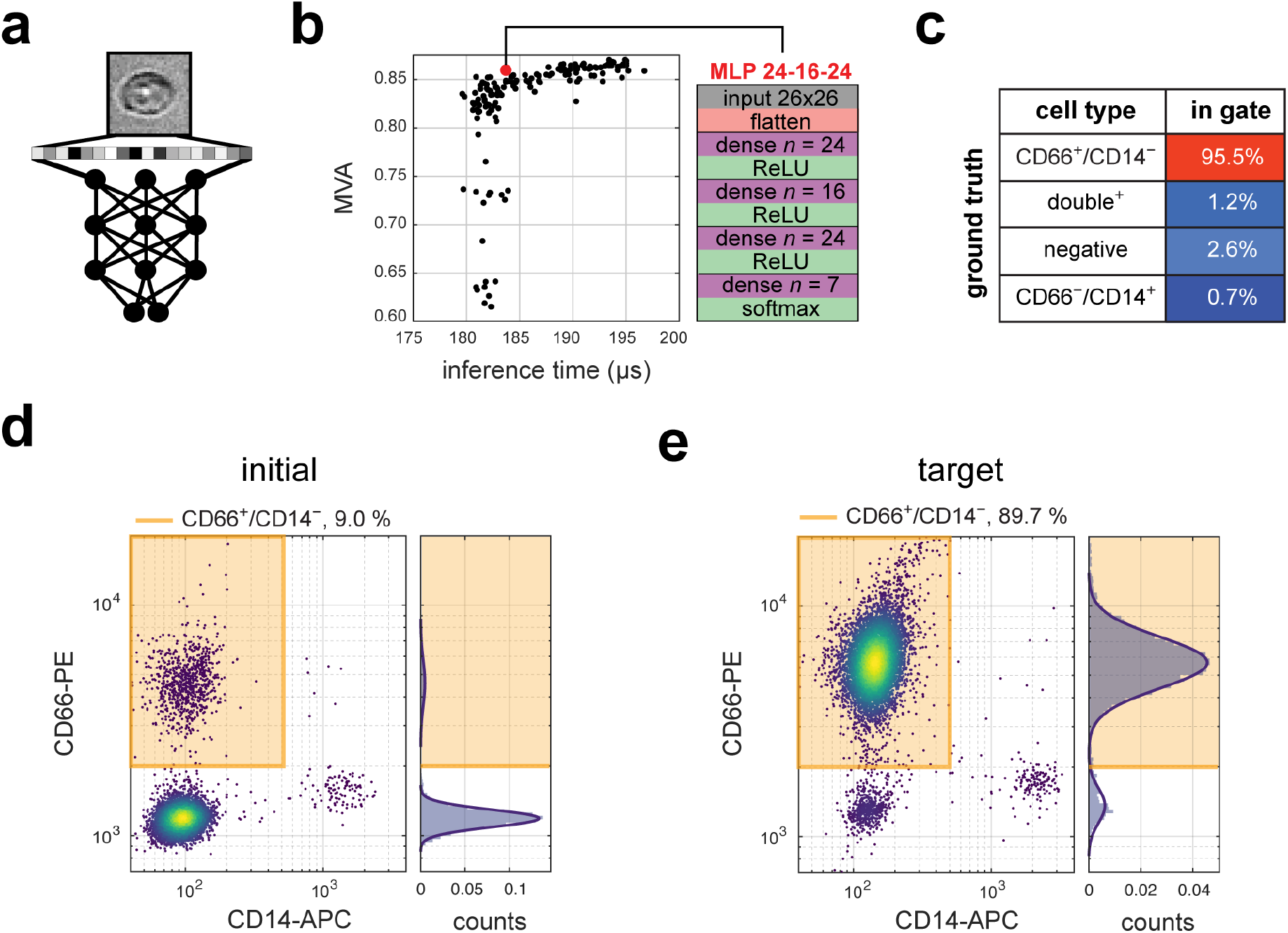
DNN-based sorting of neutrophils from RBC-depleted blood. (**a**) Schematic representation of DNN-based image analysis. (**b**) MVA versus inference time for 162 tested differently complex MLPs. The MLP selected for sorting (MLP 24-16-24) is indicated in red; its architecture details are summarized on the right. (**c**) Percentage of different cell types classified as neutrophils by DNN. Calculation based on fluorescent staining: CD66^+^/CD14^−^ cells correspond to neutrophils, CD66-/CD14+ to monocytes, double negative cells to RBCs and double positive are staining errors or cell doublets. (**d-e**) Post-analysis of the initial (**d**) and the target (**e**) samples using RT-FDC and CD14/CD66 staining.

To demonstrate how such transferring of molecular specificity into label-free DNN-based sorting can be implemented, we have used highly optimized libraries^61–63^ to train DNNs on a standard PC with the aim to distinguish neutrophils based on bright-field images alone in the RBC-depleted blood samples. First, we set out to select a neural network architecture that would be robust enough for our application. To this end, we tested the performance of 162 multilayer perceptrons (MLPs, a class of artificial neural networks) on a hand-labelled dataset, and settled on an architecture that offered a good trade-off between maximum validation accuracy (MVA = 85.9 %) and inference time (183.7 μs) (**Figure 4b**). Prior to the sorting experiment, we recorded a training set consisting of several tens of thousands of CD66/CD14-labelled blood cells images using RT-FDC. During acquisition of the training set, the focus was altered in order to account for noise that can appear during soRT-FDC experiments. After the experiments, images were cropped to 26 × 26 pixels with the cell body centered and augmented by randomly altering brightness and rotation. Experimental alterations (focus change) and computational augmentation (brightness and rotation) help to train more robust models that perform well even when applied to previously unseen cell images, which is essential for prospective sorting. CD66^+^/CD14^−^ cells were classified as neutrophils and the dataset was fed for training to the selected MLP architecture. We then applied the trained MLP on a separately recorded validation set and observed a 95.5% classification accuracy as verified by fluorescent label-based cell assignment as ground truth (**Figure 4c**). Finally, we used this DNN-classifier to sort neutrophils in realtime from RBC-depleted blood based on images alone. The post-analysis of the target as well as the initial sample was performed using RT-FDC and CD14/CD66 staining. We obtained a 89.7% purity and an 10-fold enrichment of CD66^+^/CD14^−^ cells (**Figure 4d,e**).

## Discussion

Enriching cells of interest within heterogeneous populations can be based on molecular labels^13,14,21,24^ or morpho-rheological properties such as cell size or deformability^15–19^. The latter is particularly interesting because it allows for label-free cell type purification, which is of particular value for downstream applications. Additionally, the development of machine learning approaches allows now for cell identification and sorting without prior knowledge of features distinguishing cells of interest. In this work, we introduce soRT-FDC, an on-demand label-free cell sorting approach based on RT-FDC, SSAW, and artificial intelligence, that synergizes all of the aforementioned sorting modalities. With its use we demonstrate active sorting based on fluorescence (**Figure 2d**, **Supplementary Figure 3a**) and image-derived features, such as size (**Figure 2d**, **Supplementary Figure 3b**, **Supplementary Figure 4**), deformation (**Figure 2d**, **Supplementary Figure 3a**, **Supplementary Figure 5**), and brightness (**Figure 2d**, **Supplementary Figure 3b**) of polymer bead cell mimics as well as heterogeneous cell samples, reaching purities of up to 98%. We further show that cell alignment and deformation in a narrow constriction can be beneficial for discriminating between different cell populations. Finally, we demonstrate the possibility of transferring molecular specificity of cell markers into label-free sorting of neutrophils from blood samples with the aid of DNN.

One of the most basic physical properties of cells, which can be used for sorting or for monitoring cell behavior, is its size. In conventional flow cytometry, cell size is inferred from measurements of the FSC signal intensity^24^. While this quantity can provide a good representation of the cell diameter in many cases, it is not an absolute readout of cell size since FSC depends also on features such as, for example, the refractive index of the measured object^24,27^. As demonstrated in **Supplementary Figure 2**, when measuring beads made of different materials, FSC signals fail to evaluate the relative bead sizes correctly. In our setup we rely on a direct measurement of cell size based on image-derived cell contour, which provides a physical value of cell cross-sectional area that can be converted to cell volume by contour rotation^64^, and provides a more reliable measure of object size.

Another relevant physical property of cells that can be used for sorting with soRT-FDC is cell deformation. Cell deformation, together with size, gives access to mechanical properties of cells and can, in fact, be converted to Young’s modulus in realtime using a look-up table obtained from numerical simulations^49,65^. Sorting for cell mechanics opens up versatile research venues both in basic science and in clinical research. Since the knowledge about molecular mechanisms controlling cell mechanics is still limited, sorting for the stiff or soft fraction of cells and performing genomic, transcriptomic, or proteomic analysis could foster identification of cell mechanics regulators and elucidate target molecules for on-demand modification of cell stiffness. In case of clinical applications, enrichment of small and deformable cells, which were shown to have improved capability of passing the microvasculature^40^, could aid in attaining successful delivery of cells to target organs in cell-based therapies with, for example, hematopoietic stem cells.

What distinguishes our sorting modality from other techniques for mechanics-based cell sorting is the active sorting mechanism, and the possibility to freely combine sorting for mechanics and other available parameters, such as cell size, fluorescence, or brightness. While previous studies have shown active sorting for cell mechanics^66,67^, we provide its first realization with practically useful throughput. Additionally, a range of techniques enabling passive mechanics-based sorting such as DLD^19,68^, inertial microfluidics^69^, acoustophoresis^70,71^, or filtration-based approaches^15,16^, have been developed. These techniques provide high throughput; however, they often convolve cell deformability with size — small stiff cells are sorted with large soft ones — and, since the sorting parameters are hard-wired into the device design, they do not offer flexibility in choosing sorting parameters on demand. Moreover, unlike soRT-FDC (see **Supplementary Figure 3**), the passive techniques do not provide the possibility of combining physical parameters with fluorescence readouts during sorting, which can be of use for many applications^45^.

Deformation of cells induced in the narrow channel constriction of the microfluidic chip not only enables evaluation of cell mechanical properties, but also aids in revealing differences in cell subpopulations that would be concealed otherwise. In **Figure 2** we have shown how aligning and deforming cells in the analysis channel facilitates identification of blood cell types. Another example of how deformation can aid in capturing differences between cells is the detection of plasmodium-infected RBCs in RT-FDC. While deformed, the infected RBCs are distinguishable from uninfected ones by the appearance of a characteristic feature: a white spot within the cell, likely being a vacuole^39^. This difference between infected and uninfected cells could not be picked up in static light scattering of stationary, undeformed cells in a previous study^72^.

Label-free sorting of cells, including feature-based as well as featureless Al-based sorting, is of high interest for downstream applications. In our setup it is possible to sort cells based on many image-derived features, such as cell deformation or brightness, which do not require introducing labels. Such parameters can be used for label-free identification of, for example, plasmodium-infected RBCs, or leukocytes with engulfed bacteria and fungi, which could be used for downstream genotyping. Furthermore, we demonstrate that the molecular specificity conferred by fluorescent markers can be transferred to label-free sorting with the aid of DNN for virtually any application of interest. For example, iPSC-derived cells intended for transplantation are conventionally identified with fluorescent reporters. The specificity of these reporters could be used for DNN training, and the actual sorting of unlabeled cells could then be performed exclusively based on images. This approach eliminates the need of altering the cells with markers, and avoids extra cost and lengthy preparation procedures. Additionally, since most of the tools used in current research rely on molecular markers for cell identification, they remain blind to populations for which labels are not established. Ultimately, massive amounts of images, in tandem with machine learning approaches, could be used to identify previously unknown cell populations.

While the benchmark soRT-FDC throughput of 100 cells/s is compelling, it is far from reaching the throughput of 30-50k cells/s offered by FACS. At present, the throughput of soRT-FDC is limited by the time a cell needs to be exposed to SSAW for sufficient deflection (2 ms), which, together with signal latency of 1 ms, amounts to a total cell processing time of 3 ms. To keep the number of falsely sorted cells low, only one cell should be present in the analysis and sorting regions within this time. Therefore, the concentration of used samples needs to be restricted. For faster deflection of cells, redesigning IDT geometry and using higher SSAW power could be implemented. However, higher power could lead to overheating and consequent problems with both chip integrity and cell viability. Boosting the image processing time could, in turn, be achieved by specialized hardware such as field-programmable gate arrays or GPUs.

Another technical challenge, introduced to soRT-FDC by the use of SSAW, is the decreased image quality due to scarceness of light. The lithium niobate substrate, used to induce surface acoustic wave propagation, is birefringent, which necessitates the use of a polarizer to remove the twin image. This reduces the amount of light coming through the system by half as compared to using a plain glass substrate. Even when compensating for this reduction by increased lamp brightness, the image quality is still lower than with a glass-based chip. Better images would potentially lead to improved DNN training and classification. Therefore, it could be beneficial to implement alternative sorting mechanisms, such as vapor bubble generation with microheaters^73,74^ or direct mechanical actuation of a dual-membrane system by piezoelectric elements^25,75^, which would eliminate the need for a special substrate.

Taken together, the combination of RT-FDC and SSAW-based cell sorting in conjunction with DNN classification provides a flexible sorting platform, capable of not only parameter-based sorting but also of automated image-based separation of cells. The latter provides for the opportunity to transfer molecular specificity into label-free cell sorting and to identify new cell types, the distinction of which is not possible based on known features. We anticipate broad use of soRT-FDC in both basic research and medical applications, immediate examples include label-free sorting of stem cells or retinal precursor cells for transplantation purposes.

## Acknowledgements

We thank Despina Soteriou (MPL Erlangen) for engaging discussions, and Isabel Richter (TU Dresden) and Christine Schweitzer (MPL Erlangen) for technical assistance. We further thank the Microstructure Facility at the Center for Molecular and Cellular Bioengineering (CMCB) at Technische Universität Dresden (in part funded by the State of Saxony and the European Regional Development Fund) for hosting the chip fabrication. The HL60/S4 cells were a kind gift of D. and A. Olins (University of New England), Kc167 cells were a kind gift of B. Baum (University College London). The authors acknowledge funding from the Alexander von Humboldt-Stiftung (Alexander von Humboldt Professorship to J.G.), ERC Starting Grant (starting grant “LightTouch” #282060 to J.G.), European Commission through LAPASO ITN project (EU FP7/2007-2013, #607350), a DKMS ‘Mechthild Harf Research Grant’ (DKMS-SLS-MHG-2016-02 to A.J.), and German Research Foundation (#GU 612/5-1 to J.G.).

## Author contributions

J.G. conceived the project. A.A.N. developed and built the microfluidic chips and SSAW actuators. M.N. P.R. and A.A.N. integrated SSAW sorting into the experimental setup. M.N., with the contributions of P.R. and C.H., developed the software for real-time sorting. S.A. performed flow simulations validating chip design. A.A.N., with the contributions of M.H., M.Kr., A.J., P.R., M.U., N.T. and M.Ku., performed the sorting experiments with beads, cells, and blood samples. M.Kr., A.J., M.H. and N.T. prepared the blood samples and developed cell staining protocols. M.H. developed the DNN analysis and performed the AI sorting experiments. S.G. and R.G. produced the polyacrylamide beads. S.G., R.G. and F.R. provided advice and technical support for chip production. A.T. performed AFM measurements. M.U., M.H. and A.A.N. analyzed the data. P.M. developed *ShapeOut* analysis software. M.U. visualized the data and prepared figures. M.U., with the support of J.G., P.R., M.H. and A.A.N. prepared the initial version of the manuscript. All authors revised and edited the manuscript. J.G. and A.J. acquired funding.

## Data availability statement

The data acquired in the experiments presented in this study are available on figshare at http://doi.org/10.6084/m9.figshare.11302595.v1.

## Code availability

*ShapeOut* and *AID* are open source and available on GitHub at URLs provided in **Methods**. *Shape1n* is commercially available from Zellmechanik Dresden GmbH. The C++ code used for sorting is custom-built for the hardware used in the setup and available onsite to interested parties.

## Methods

### soRT-FDC chip design

Schematic illustrations of the soRT-FDC chip layout are presented in **Figures 1a** and **2a**. The sheath and sample inlet regions follow a previously published layout^41,45^, with an additional serpentine-like curved channel in the sample inlet region (see **Figure 2a**) which aids in focusing and longitudinal ordering of cells^76^. Upon contact with sheath fluid, cells enter a 20-μm or 30-μm wide and 880-μm long channel, in which cell deformation is induced and cell images, as well as fluorescence signals, are recorded. Afterwards, the channel broadens to 50 μm for a 200-μm long stretch where SSAW are applied by IDTs located on the channel sides. This 50-μm wide region is followed by a bifurcation dividing the flow into a target and default branch. Finite element flow simulations in the channel were conducted using COMSOL (COMSOL Multiphysics, Sweden) to analyze the velocity fields and verify the bifurcation position. The bifurcation was positioned 5 μm off-center to ensure that all cells move towards the default outlet if the SSAW are switched off (see **Supplementary Figure 1**).

### Chip fabrication and assembly

A soRT-FDC chip consists of a polydimethylsiloxane (PDMS) replica bonded to a lithium niobate substrate with chromium-gold IDTs deposited on its surface. PDMS replicas were fabricated according to well established soft-lithography processes. In brief, 20-μm or 30-μm high layer of AZ 15nXT (450 CPS) photoresist (MicroChemicals, Germany) was spun onto a 4” silicon wafer using a spin coater (Laurell WS-650Mz-23NPP, Laurell Technologies, PA, USA) and a master mold was developed using UV-exposure. 1H,1H,2H,2H-perfluorooctyl-trichlorosilane (#448931, Sigma-Aldrich, Germany) vapor was deposited onto the developed master mold under vacuum to provide hydrophobic coating and ease the peel-off of the replicas. PDMS (base to curing agent ratio of 10:1 w/w; #634165S, SYLGARD 184, VWR, Germany) was poured over the master, degassed, and cured at 65°C for six hours. The fabrication of IDTs onto 128° Y-cut lithium niobate (LiNbO3, Roditi International, UK) substrate was performed via photolithography on a thin layer of AZ 5214E image reversal resist (MicroChemicals, Germany), metal evaporation of chromium and gold layers (Cr/Au, 10 nm/70 nm, respectively; Kurt J. Lesker, UK) in a thermal evaporation system (NANO36; Kurt J Lesker, UK) and subsequent lift-off. Each IDT has 40 electrode pairs, an aperture of 200 μm, and an inter-finger distance of 70 μm, resulting in an excitation frequency of 55.23 MHz. The resonance frequency SSAW return loss (S11) = −2.8 dBm. An additional 100 nm layer of SiO2 was deposited on top of the substrate, which drastically improved the PDMS bonding and permitted the use of much higher flow rates without leakage. Before bonding, holes were punched into the PDMS slab at inlet and outlet positions using 1.5-mm biopsy punchers (#49115, pfm medical, Germany). The substrate and the PDMS were covalently bonded to prevent leakage using plasma activation (50 W, 30 s; Plasma Cleaner Atto; Diener electronic, Germany). The bonded devices were cured in an oven at 65°C for 48 hours. The sheath and sample fluid syringes (BD Luer-Lok™ 5-ml syringe #613-2043P and BD Luer-Lok™ 1-ml syringe #613-4971; VWR, Germany) were connected to the fabricated chip via FEP Tubing (1/16” OD, 0.03” ID; #1520XL, Postnova Analytics, Germany). The fluids were flowed through the chip using high-precision syringe pump (three modules, neMESyS 290N, NeMESyS, Cetoni, Germany). The default outlet was connected to a 1-ml syringe and fitted into the third pump module which was operated in withdrawal mode. This additional withdrawal helps to avoid accidental slipping of cells to the target outlet. The target outlet was fitted with tubing which led into a collection vial. For the 20-μm chip, flow rates of 0.01, 0.03 and ca. −0.027 μl/s were used for the sample, sheath, and default modules, respectively. For the 30-μm chip flow rates of 0.02, 0.06, and ca. −0.05 μl/s were applied.

### SSAW sorting mechanism

Two IDTs were placed at opposing channel sides and actuated at their resonance frequency, *f*, given by:

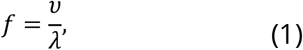

where *ν* is the velocity of sound in the lithium niobate substrate (3,978 m s^-1^) and *λ* is the acoustic wavelength, defined by the distance between adjacent IDT fingers. The constructive interference of counter-propagating waves generated by opposing IDTs results in the emergence of standing surface acoustic waves (SSAW). The SSAW were tuned to have one pressure node positioned in front of the target outlet. SSAW generated an acoustic radiation force, *F_r_*, that pushed cells towards the pressure node and directed them into the target outlet. *F_r_* is given by the following formula^77^:

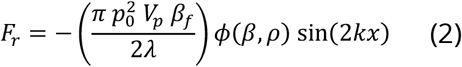

where *p_0_* is the acoustic pressure, *λ* is the acoustic wavelength, *V_p_* is the volume of the particle, *β* is compressibility, *ρ* is density, and *φ* is the acoustic contrast factor that defines if the particle will translate to the pressure node (*φ* >0) or the pressure antinode (*φ* <0) defined as:

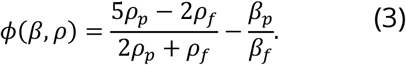

The subscripts *p* and *f* stand for particle and fluid, respectively.

### Experimental setup and sorting feedback system

The experiments were performed using an RT-FDC setup^45^ with a digital output (TTL) controlling the actuation of SSAW function generator (BSG F20, BelektroniG, Germany). The basic RT-FDC setup consists of an inverted microscope (Axio Observer Z1, Zeiss, Germany) and a pulsed, high-power controlled by a custom driver circuit (AcCellerator L1, Zellmechanik Dresden, Germany). Images captured by a highspeed CMOS camera (EoSens CL, MC1362, Mikrotron, Germany) were transferred to a standard PC using a full camera-link frame grabber card (PCIe-1433, National Instruments, TX, USA). The camera operates at 2,500 frames per second and triggers short LED light pulses (2 μs) to prevent image blurring. Real-time image processing is performed by a custom C++ based sorting software, which leverages the OpenCV computer vision library^78^, returning a set of parameters describing the captured object (summarized in **Supplementary Table 1**). If the values of these parameters fall into a user-specified sorting gate, a trigger signal from the frame grabber card is induced to the SSAW function generator. The delay induced by image processing is on average 225 μs. The total delay between image exposure and sorting trigger was measured to be 1 ms. During sorting experiments using a 20-μm analysis channel, we apply a constant flow rate of 0.04 μl/s, resulting in an object speed of approximately 20 cm/s. Based on this speed and the trigger delay, the cell travels ca. 120 μm before the SSAW can switch on, therefore the measurement region of interest is placed 120 μm before the SSAW sorting region. In the case of 30-μm analysis channel used with a flow rate of 0.08 μl/s, the measurement region of interest is placed 180 μm before the SSAW sorting region. For imaging during sorting, a 20× objective (Plan-Apochromat, 20×/0.8; #440640-9903, Zeiss, Germany) and additional polarizer (Polarizer D, 90° rotatable, removable; #427706-0000-000, Zeiss) were used. For most of the post-analysis measurements, standard RT-FDC chips^45^ with glass coverslip bottom, in combination with a 20× objective or a 40× objective (EC-Plan-Neofluar, 40×/0.75; #420360-9900, Zeiss, Germany), were used. For details of experimental conditions used in every measurement see **Supplementary Table 3**.

### Data acquisition and analysis

For the acquisition of data during sorting a custom C++ program, that also executes sorting, was used. For the acquisition of post-analysis measurements, an RT-FDC setup^45^ and *ShapeIn* software (*ShapeIn2*; Zellmechanik Dresden, Germany) were used. The sizes of acquired images varied between 100 × 40 pixels and 250 × 100 pixels, depending on the width of the measurement channel and objective magnification used, and are listed in **Supplementary Table 4**. The data was postprocessed using *ShapeOut (ShapeOut 0.9.5;* https://github.com/ZELLMECHANIK-DRESDEN/ShapeOut; Zellmechanik Dresden) and *MATLAB* (MATLAB R2018b, The MathWorks, MA, United States). Details regarding filtering for plotting and purity calculations are summarized in the **Supplementary Table 5**.

### Measurement buffer

For soRT-FDC experiments and post-analysis measurements we used a measurement buffer (MB) with high viscosity, which provides for scalable cell deformation and abates cell sedimentation. MB is PBS-based solution containing 0.6 % (w/v) methyl cellulose (4000 cPs; Alfa Aesar, Germany) with pH 7.4, osmolality of 310-315 mOsm/kg and viscosity adjusted to 25 mPa s at 24°C with a falling sphere viscometer (HAAKE™ Faling Ball Viscometer Type C, Thermo Fisher Scientific, MA, USA), which corresponds to 7.3 mPa s in the measurement channel^79^. MB was used as sheath fluid and as medium for suspensions of cells or beads.

### Commercially-available beads

For the validation of size estimation (**Supplementary Figure 2** and **3b**) and sorting based on size (**Figure 1d**, middle panel) we used mixtures of several commercially-available microspheres, including melamine beads (MF-FluoBlau-L948, 9.78 ± 0.15 μm diameter), silica beads (SiO_2_-F-L3519-1, 13.79 ± 0.59 μm diameter), polystyrene beads (PS/Q-F-KM194, 15.21 ± 0.31 μm diameter), and poly(methyl methacrylate) beads (PMMM-F-B1423, 17.23 ± 0.24 μm diameter), all purchased from Microparticles, Germany.

### Polyacrylamide bead production

Polyacrylamide hydrogel beads were produced in house using PDMS-based flow-focusing microfluidic device as described elsewhere^46^. For the preparation of beads with different stiffnesses (**Figure 1d**, lower panel, **Supplementary Figure 3a**) total monomer concentrations (molar ratio of bis-acrylamide to acrylamide 1:61.5; #146072 and #A3553, Sigma-Aldrich) of 7.9%, 9.9% and 13.8% were used. To enable modification with the fluorescent dye (**Figure 1d**, upper panel, **Supplementary Figure 3a**), N-hydroxysuccinimide ester (#130672, Sigma-Aldrich) was incorporated into polyacrylamide droplets during the production. After the production process, 1 pg of AlexaFluor 488 Hydrazide (#A10436, Thermo Fisher Scientific) per bead was added.

### Cell lines and cell culture

Kc167 *Drosophila* cells (received from Buzz Baum, MRC Laboratory for Molecular Cell Biology, University College London, London, United Kingdom) were cultured at room temperature in M3 Shields and Sang medium (Invitrogen, CA, USA) supplemented with 10% heat-inactivated fetal bovine serum (FBS) (#10270106, Thermo Fisher Scientific) and 1% penicillin/streptavidin (#15140122, Thermo Fisher Scientific) at 24°C under atmospheric CO2 concentration. HL60/S4 cells (received from Donald E. Olins and Ada L. Olins, University of New England, ME, USA) were grown in ATCC-modified RPMI-1640 medium (#A1049101, Thermo Fisher Scientific) supplemented with 10% heat-inactivated FBS and 1% penicillin/streptavidin at 37°C and 5% CO2 with subculturing every second day. For experiments, cells were harvested by spinning down at 114 RCF for 5 min and re-suspended in MB at a final concentration of 1 × 10^6^ cells/ml.

### Blood sample preparation

Whole blood was drawn from healthy human donors after informed consent (ethical approval EK89032013, granted by the ethics committee of the Technische Universität Dresden) into 10 ml sodium-citrate tubes (S-Monovette^®^ 10ml 9NC; #02.1067.001, Sarstedt, Germany) and depleted for RBCs using dextran sedimentation. To this end, 6% dextran solution was prepared by dissolving dextran (Dextran T500, Pharmacosmos, Denmark) in 0.9% sodium chloride solution (0.9% Sodium Chloride Irrigation, Baxter Healthcare, Switzerland). So-prepared dextran solution was added to whole blood in a 1:4 ratio and RBCs were allowed to sediment for 30 min, forming a red pellet. RBC-depleted supernatant was collected and centrifuged for 10 min at 120 g (Universal 30RF, Hettich, Switzerland). The supernatant was discarded, and the cell pellet was resuspended in 2 ml of MB (0.6% MC). For evaluation of CD14/CD66 surface marker expression, the cells were stained with APC-conjugated anti-human CD14 (dilution 1:20, #17-0149-42, eBioscience, CA, USA) and PE-conjugated anti-human CD66a/c/e (dilution 1:40, #34303, BioLegend, CA, USA) in MB for 25 min at 600 rpm in a block thermomixer (Thermomixer comfort 5355, Eppendorf, Germany). For sorting of RBCs based on deformation the blood was used directly, without performing the dextran sedimentation step. For sorting of RBCs based on deformation, the blood was used directly, without performing the dextran sedimentation step.

### MLP Selection

For the selection of a robust MLP architecture, we used an extensive dataset of 100 measurements of RBC-depleted blood samples from 20 donors captured at 20× magnification in a soRT-FDC chip. The dataset was labelled manually with 7 categories using polygon gates in the deformation-size space to distinguish between debris, lymphocytes, RBCs, granulo-monocytes, and cell doublets. Granulo-monocytes were further classified into eosinophils, neutrophils, and monocytes using polygon gates in the brightness vs. standard deviation of brightness space. The number of all cells classified into individual cell types is listed in **Supplementary Table 6**. MLP screening was performed using a collection of models in *AID* software (*AIDeveloper_0.0.4;* https://github.com/maikherbig/AIDeveloper)^80^.

### Training of MLP for neutrophil sorting

Prior to the sorting experiment, a dataset of approximately 30,000 cells labeled with CD14/CD66 fluorescent markers was recorded fed into AID software^80^ to train the selected MLP architecture. To provide for robust training, the focus was varied during the dataset acquisition and computational augmentation of recorded images was performed by altering brightness using a linear transfer function, adding Gaussian noise, and random rotation. For the purpose of training, CD66^+^/CD14^−^ cells were classified as neutrophils. For validation, an additional dataset of approximately 10,000 cells was acquired (see **Supplementary Table 7** for a number of cells in each class). The trained DNN model was exported from AID as *\*.nnet* file using *keras2cpp* library (https://github.com/pplonski/keras2cpp). The *keras2cpp* library was then used to load the *\*.nnet* file into the C++ sorting software.

### Cell viability and proliferation assay

HL60/S4 cells were suspended in MB at 1 × 10^6^ cells/ml and exposed to SSAW during sorting in a soRT-FDC chip. The initial sample (prepared as for the measurement, but not run through the setup), as well as cells collected in the target outlet after exposure to SSAW and default outlet when SSAW was off, were collected by centrifugation at 114 RCF for 5 min and suspended in 1 ml of fresh medium in a cell culture-treated 12-well plate (#665180, Greiner Bio-One, Austria). The starting cell count was equal to 70,400 for target sample, and 100,000 for both initial and default samples. Cell count and cell viability were assessed using an automated cell counter (Countess II, AMQAX1000, Thermo Fischer Scientific) and trypan blue staining (Trypan Blue stain 0.4%, #T10282, Thermo Fischer Scientific) on the day of seeding (day 0) and on the four following days (day 1 – 4). On day 2, cells were collected by centrifugation at 114 RCF for 5 min and transferred into 2 ml of fresh medium in a cell culture-treated 6-well plate (#657160, Greiner Bio-One) to provide for sufficient space for further growth.

### AFM indentation measurements

To assess the influence of surface acoustic waves and/or circulation through the soRT-FDC microfluidic system on cell stiffness, cells were flowed through the microfluidic channel of the soRT-FDC chip either with or without SSAW-induced sorting. The initial sample (prepared as for the measurement and kept at room temperature for comparable time but not run through the setup), as well as cells collected in the target outlet after exposure to SSAW and default outlet when SSAW was off, were collected by centrifugation for 3 min at 180 RCF and seeded onto glass bottom dishes (fluorodish; #FD35100, WPI, FL, USA) in CO2-independent medium (#18045-054, Thermo Fisher Scientific). After settling down for 10 minutes and attachment to the surface, cells were mechanically probed on a Nanowizard 4 (JPK Instruments, Germany) equipped with a Petri dish heater (JPK Instruments) at 37°C. Cells were indented using an arrow-T1 (Nanoworld, Switzerland) cantilever equipped with a *R* = 2.5 μm polystyrene bead (PS-R-5.0; Microparticles) at a speed of 5 μm/s. Apparent Young’s moduli were extracted after fitting the force-indentation curves up to a maximum depth of 2 μm using the Sneddon-Hertz model for a spherical indenter in the JPK data processing software (JPK Instruments). The obtained Young’s modulus values were corrected using the simplified double-contact model accounting for the compression of cells from the bottom^81^. Datasets were compared using a Kruskal-Wallis test with a Dunn’s multiple comparisons test (GraphPad Prism; GraphPad Software, CA, USA).

## References

1. Dainiak, M. B., Kumar, A., Galaev, I. Y. & Mattiasson, B. Methods in Cell Separations. in Cell Separation 1–18 (Springer, 2007).

2. Wyatt Shields Iv, C., Reyes, C. D. & López, G. P. Microfluidic cell sorting: A review of the advances in the separation of cells from debulking to rare cell isolation. Lab Chip 15, 1230–1249 (2015).

3. Cossarizza, A. et al. Guidelines for the use of flow cytometry and cell sorting in immunological studies. Eur. J. Immunol. 47, 1584–1797 (2017).

4. Heath, J. R., Ribas, A. & Mischel, P. S. Singlecell analysis tools for drug discovery and development. Nat. Rev. Drug Discov. 15, 204–216 (2016).

5. Tanay, A. & Regev, A. Scaling single-cell genomics from phenomenology to mechanism. Nature 541, 331–338 (2017).

6. Islam, M. et al. Microfluidic cell sorting by stiffness to examine heterogenic responses of cancer cells to chemotherapy. Cell Death Dis. 9, 239 (2018).

7. Poser, S. W. et al. Controlling distinct signaling states in cultured cancer cells provides a new platform for drug discovery. FASEB J.. 33, 9235–9249 (2019).

8. Handgretinger, R. et al. Isolation and transplantation of autologous peripheral CD34+ progenitor cells highly purified by magnetic-activated cell sorting. Bone Marrow Transplant. 21, 987–993 (1998).

9. Stamm, C. et al. Autologous bone-marrow stem-cell transplantation for myocardial regeneration. Lancet 361, 45–46 (2003).

10. Bartsch, U. et al. Retinal cells integrate into the outer nuclear layer and differentiate into mature photoreceptors after subretinal transplantation into adult mice. Exp. Eye Res. 86, 691–700 (2008).

11. Gagliardi, G. et al. Characterization and Transplantation of CD73-Positive Photoreceptors Isolated from Human iPSC-Derived Retinal Organoids. Stem Cell Reports 11, 665–680 (2018).

12. Majzner, R. G. & Mackall, C. L. Clinical lessons learned from the first leg of the CAR T cell journey. Nat. Med. 25, 1341–1355 (2019).

13. Miltenyi, S., Müller, W., Weichel, W. & Radbruch, A. High gradient magnetic cell separation with MACS. Cytometry 11, 231–238 (1990).

14. Šafařik, I. & Šafařiková, M. Use of magnetic techniques for the isolation of cells. J. Chromatogr. B Biomed. Sci. Appl. 722, 33–53 (1999).

15. Preira, P. et al. Passive circulating cell sorting by deformability using a microfluidic gradual filter. Lab Chip 13, 161–170 (2013).

16. Wang, G. et al. Microfluidic cellular enrichment and separation through differences in viscoelastic deformation. Lab Chip 15, 532–540 (2015).

17. Cheng, Y., Ye, X., Ma, Z., Xie, S. & Wang, W. High-throughput and clogging-free microfluidic filtration platform for on-chip cell separation from undiluted whole blood. Biomicrofluidics 10, (2016).

18. Huang, L. R., Cox, E. C., Austin, R. H. & Sturm, J. C. Continuous Particle Separation Through Deterministic Lateral Displacement. Science (80-.). 304, 987–990 (2004).

19. Beech, J. P., Holm, S. H., Adolfsson, K. & Tegenfeldt, J. O. Sorting cells by size, shape and deformability. Lab Chip 12, 1048–1051 (2012).

20. MacDonald, M. P., Spalding, G. C. & Dholakia, K. Microfluidic sorting in an optical lattice. Nature 426, 421–424 (2003).

21. Bonner, W. A., Hulett, H. R., Sweet, R. G. & Herzenberg, L. A. Fluorescence Activated Cell Sorting. Rev. Sci. Instrum. 43, 404–409 (1972).

22. Schmid, L., Weitz, D. A. & Franke, T. Sorting drops and cells with acoustics: Acoustic microfluidic fluorescence-activated cell sorter. Lab Chip 14, 3710–3718 (2014).

23. Nawaz, A. A. et al. Acoustofluidic Fluorescence Activated Cell Sorter. Anal. Chem. 87, 12051–12058 (2015).

24. Shapiro, M. H. Practical Flow Cytometry. (John Wiley & Sons, Inc., 2003). doi:10.1002/0471722731.

25. Nitta, N. et al. Intelligent Image-Activated Cell Sorting. Cell 175, 266–276.e13 (2018).

26. Ota, S. et al. Ghost cytometry. Science (80-.). 360, 1246–1251 (2018).

27. Salzman, G. C. Light Scattering Analysis of Single Cells. in Cell Analysis 111–143 (Springer US, 1982). doi:10.1007/978-1-4684-4097-3_5.

28. Hiramatsu, K. et al. High-throughput label-free molecular fingerprinting flow cytometry. Sci. Adv. 5, 1–9 (2019).

29. Suzuki, Y. et al. Label-free chemical imaging flow cytometry by high-speed multicolor stimulated Raman scattering. Proc. Natl. Acad. Sci. 116, 15842–15848 (2019).

30. Gorthi, S. S. & Schonbrun, E. Phase imaging flow cytometry using a focus-stack collecting microscope. Opt. Lett. 37, 707 (2012).

31. Chen, C. L. et al. Deep Learning in Label-free Cell Classification. Sci. Rep. 6, 1–16 (2016).

32. Merola, F. et al. Tomographic flow cytometry by digital holography. Light Sci. Appl. 6, 1–7 (2017).

33. Min, J. et al. Quantitative phase imaging of cells in a flow cytometry arrangement utilizing Michelson interferometer-based off-axis digital holographic microscopy. J. Biophotonics 1–10 (2019) doi:10.1002/jbio.201900085.

34. Zhang, J., Nou, X. A., Kim, H. & Scarcelli, G. Brillouin flow cytometry for label-free mechanical phenotyping of the nucleus. Lab Chip 17, 663–670 (2017).

35. Zhang, J., Fiore, A., Yun, S. H., Kim, H. & Scarcelli, G. Line-scanning Brillouin microscopy for rapid non-invasive mechanical imaging. Sci. Rep. 6, 1–8 (2016).

36. Urbanska, M. et al. A comparison of microfluidic methods for high throughput cell deformability measurements. Nat. Methods (in press).

37. Tavares, S. et al. Actin stress fiber organization promotes cell stiffening and proliferation of pre-invasive breast cancer cells. Nat. Commun. 8, (2017).

38. Kräter, M. et al. Alterations in Cell Mechanics by Actin Cytoskeletal Changes Correlate with Strain-Specific Rubella Virus Phenotypes for Cell Migration and Induction of Apoptosis. Cells 7, 136 (2018).

39. Toepfner, N. et al. Detection of human disease conditions by single-cell morpho-rheological phenotyping of blood. Elife 7, e29213 (2018).

40. Tietze, S. et al. Spheroid Culture of Mesenchymal Stromal Cells Results in Morphorheological Properties Appropriate for Improved Microcirculation. Adv. Sci. 1802104 (2019) doi:10.1002/advs.201802104.

41. Otto, O. et al. Real-time deformability cytometry: on-the-fly cell mechanical phenotyping. Nat. Methods 12, 199–202 (2015).

42. Basiji, D. A., Ortyn, W. E., Liang, L., Venkatachalam, V. & Morrissey, P. Cellular Image Analysis and Imaging by Flow Cytometry. Clin. Lab. Med. 27, 653–670 (2007).

43. Blasi, T. et al. Label-free cell cycle analysis for high-throughput imaging flow cytometry. Nat. Commun. 7, 10256 (2016).

44. Doan, M. et al. Diagnostic Potential of Imaging Flow Cytometry. Trends Biotechnol. 36, 649–652 (2018).

45. Rosendahl, P. et al. Real-time fluorescence and deformability cytometry. Nat. Methods 15, 355 (2018).

46. Girardo, S. et al. Standardized microgel beads as elastic cell mechanical probes. J. Mater. Chem. B 6, 6245–6261 (2018).

47. Marshall, W. F. et al. The big question of cell size. BMC Biol. 10, 101 (2012).

48. Mietke, A. et al. Extracting Cell Stiffness from Real-Time Deformability Cytometry: Theory and Experiment. Biophys. J. 109, 2023–2036 (2015).

49. Mokbel, M. et al. Numerical Simulation of Real-Time Deformability Cytometry To Extract Cell Mechanical Properties. ACS Biomater. Sci. Eng. 3, 2962–2913 (2017).

50. Hartono, D. et al. On-chip measurements of cell compressibility via acoustic radiation. Lab Chip 11, 4072–4080 (2011).

51. Gustafson, M. P. et al. A method for identification and analysis of nonoverlapping myeloid immunophenotypes in humans. PLoS One 10, e0121546 (2015).

52. Bashant, K. R. et al. Real-time deformability cytometry reveals sequential contraction and expansion during neutrophil priming. J. Leukoc. Biol. 105, 1143–1153 (2019).

53. Das, D., Ghosh, M., Chakraborty, C., Maiti, A. K. & Pal, M. Probabilistic prediction of malaria using morphological and textural information. in 2011 International Conference on Image Information Processing 1–6 (IEEE, 2011). doi:10.1109/ICIIP.2011.6108879.

54. Linder, N. et al. A malaria diagnostic tool based on computer vision screening and visualization of Plasmodium falciparum candidate areas in digitized blood smears. PLoS One 9, e104855 (2014).

55. Adjed, F., Faye, I., Ababsa, F., Gardezi, S. J. & Dass, S. C. Classification of skin cancer images using local binary pattern and SVM classifier. in 080006 (2016). doi:10.1063/1.4968145.

56. Hamilton, N. A., Pantelic, R. S., Hanson, K. & Teasdale, R. D. Fast automated cell phenotype image classification. BMC Bioinformatics 8, 110 (2007).

57. Sarrafzadeh, O., Dehnavi, A. M., Banaem, H. Y., Talebi, A. & Gharibi, A. The Best Texture Features for Leukocytes Recognition. J. Med. Signals Sens. 7, 220–227.

58. Ge, Y. et al. Cell Mechanics Based Computational Classification of Red Blood Cells Via Unsupervised Machine Intelligence Applied to Morpho-Rheological Markers. IEEE IEEE/ACM Trans. Comput. Biol. Bioinforma.

59. Vandenberghe, M. E. et al. Relevance of deep learning to facilitate the diagnosis of HER2 status in breast cancer. Sci. Rep. 7, 45938 (2017).

60. Buggenthin, F. et al. Prospective identification of hematopoietic lineage choice by deep learning. Nat. Methods 14, 403–406 (2017).

61. Nickolls, J., Buck, I., Garland, M. & Skadron, K. Scalable parallel programming with CUDA. Queue 6, 40 (2008).

62. Abadi, M. et al. TensorFlow: Large-Scale Machine Learning on Heterogeneous Distributed Systems. Preprint at https://arxiv.org/abs/1603.04467. (2016).

63. Al-Rfou, R. et al. Theano: A Python framework for fast computation of mathematical expressions. Preprint at https://arxiv.org/abs/1605.02688. (2016).

64. Herbig, M., Mietke, A., Müller, P. & Otto, O. Statistics for real-time deformability cytometry: Clustering, dimensionality reduction, and significance testing. Biomicrofluidics 12, 042214 (2018).

65. Urbanska, M., Rosendahl, P., Kräter, M. & Guck, J. High-throughput single-cell mechanical phenotyping with real-time deformability cytometry. Methods Cell Biol. 147, 175–198 (2018).

66. Yang, T. et al. An integrated optofluidic device for single-cell sorting driven by mechanical properties. Lab Chip 15, 1262–1266 (2015).

67. Faigle, C. et al. A monolithic glass chip for active single-cell sorting based on mechanical phenotyping. Lab Chip 15, 1267–1275 (2015).

68. Holmes, D. et al. Separation of blood cells with differing deformability using deterministic lateral displacement. Interface Focus 4, 20140011 (2014).

69. Di Carlo, D., Edd, J. F., Irimia, D., Tompkins, R. G. & Toner, M. Equilibrium separation and filtration of particles using differential inertial focusing. Anal. Chem. 80, 2204–2211 (2008).

70. Ding, X. et al. Cell separation using tilted-angle standing surface acoustic waves. Proc. Natl. Acad. Sci. U. S. A. 111, 12992–12997 (2014).

71. Augustsson, P., Karlsen, J. T., Su, H. W., Bruus, H. & Voldman, J. Iso-acoustic focusing of cells for size-insensitive acousto-mechanical phenotyping. Nat. Commun. 7, 1–9 (2016).

72. Park, Y. et al. Static and dynamic light scattering of healthy and malaria-parasite invaded red blood cells. J. Biomed. Opt. 15, 020506 (2010).

73. Pritchard, R. H. et al. Cell sorting actuated by a microfluidic inertial vortex. Lab Chip 19, 2456–2465 (2019).

74. De Wijs, K. et al. Micro vapor bubble jet flow for safe and high-rate fluorescence-activated cell sorting. Lab Chip 17, 1287–1296 (2017).

75. Sakuma, S., Kasai, Y., Hayakawa, T. & Arai, F. On-chip cell sorting by high-speed local-flow control using dual membrane pumps. Lab Chip 17, 2760–2767 (2017).

76. Di Carlo, D., Irimia, D., Tompkins, R. G. & Toner, M. Continuous inertial focusing, ordering, and separation of particles in microchannels. Proc. Natl. Acad. Sci. U. S. A. 104, 18892–18897 (2007).

77. Ding, X. et al. Surface acoustic wave microfluidics. Lab Chip 13, 3626–3649 (2013).

78. Bradski, G. The OpenCV library. Dr Dobbs J. Softw. Tools 25, 120–126 (2000).

79. Herold, C. Mapping of Deformation to Apparent Young’s Modulus in Real-Time Deformability Cytometry. Preprint at https://arxiv.org/abs/1704.00572. (2017).

80. Kräter, M. et al. AI Developer: deep learning image classification in life science and beyond. Preprint at https://www.biorxiv.org/content/10.1101/2020.03.03.975250v1. (2020).

81. Glaubitz, M. et al. A novel contact model for AFM indentation experiments on soft spherical cell-like particles. Soft Matter 10, 6732–6741 (2014).

