## Supplementary Information for "Using real-time fluorescence and deformability cytometry and deep learning to transfer molecular specificity to label-free sorting"

**Supplementary Table 1. Features available in real-time that can be used for sorting.**

| feature name | description |
|---|---|
| **supported for AI and non-AI sorting** | |
| length (a) | size of the object in the direction parallel to the flow |
| height (b) | size of the object in the direction perpendicular to the flow |
| aspect ratio (a/b) | ratio between object's length and height |
| **supported for non-AI sorting** | |
| area raw | area enclosed by the contour fitted to the object |
| area (size) | area enclosed by the convex hull of the contour fitted to the object |
| area ratio | ratio between area raw and area |
| deformation | $1 - \frac{2\sqrt{\pi\, area}}{perimeter}$ |
| inertia ratio | $\frac{I_{yy}}{I_{xx}}$ <br> $I_{yy}$ - second moment of contour area calculated for $y$-direction <br> $I_{xx}$ - second moment of contour area calculated for $x$-direction <br> $I_{yy} = \iint_A y^2 dx\, dy$, $I_{xx} = \iint_A x^2 dx\, dy$ <br> $x$ and $y$ represent Cartesian coordinates |
| Young's modulus | mechanical property that quantifies cell stiffness derived from numerical simulations[49], in real-time obtained from a look-up table based on object size and deformation |
| brightness | average brightness of pixels enclosed by the contour fitted to the object |
| s.d. brightness | standard deviation of the brightness of pixels enclosed by the contour fitted to the object |
| FL-1 intensity | maximum fluorescence intensity recorded in channel 1; excitation laser $\lambda$ = 488 nm, emission filter 525/50 |
| FL-2 intensity | maximum fluorescence intensity recorded in channel 2; excitation laser $\lambda$ = 561 nm, emission filter 593/46 |
| FL-3 intensity | maximum fluorescence intensity recorded in channel 3; excitation laser $\lambda$ = 640 nm, emission filter 700/75 |

**Supplementary Table 2| Percentages of different cell subpopulations in initial sample, as well as in the samples after sorting for red blood cells (RBC), lymphocytes (ly), and myeloid cells (my).** The percentages were calculated based on 2D gates in cell size-deformation space as indicated in **Figure 2**.

| sample | % of all cells | | | | fold enrichment |
|---|---|---|---|---|---|
| | **RBC** | **ly** | **my** | **rest** | |
| initial | 49.5 | 4.1 | 43.2 | 3.2 | - |
| sorted RBC | 96.6 | 0.1 | 1.5 | 1.9 | 1.95 × |
| sorted ly | 3.2 | 89.1 | 4.4 | 3.3 | 21.7 × |
| sorted my | 1.6 | 0.6 | 94.7 | 3.1 | 2.0 × |

**Supplementary Table 3| Experimental details for each of the soRT-FDC measurements presented.** The details include the size of the analysis channel ($s_{ch}$), the power of lasers used during data acquisition and/or post-analysis, the substrate of the chip as well as the objective magnification used during post-analysis, and the summary of gates used during sorting. PAA – polyacrylamide, def – deformation, br – brightness, $LiNbO_3$ – lithium niobate.

| | | | laser settings | | post-analysis | | sorting gates | | | | |
|---|---|---|---|---|---|---|---|---|---|---|---|
| | **sample** | **$s_{ch}$ (μm)** | **sort** | **post-analysis** | **chip subst.** | **objec tive** | **size (μm$^2$)** | **def** | **br (a.u.)** | **FL-1 (a.u.)** | **area ratio** |
| **Figures** | | | | | | | | | | | |
| **1d** | PAA beads, FL-1 sort | 20 | FL-1: 35% | FL-1: 35% | glass | 20x | x | x | x | 1000 – 10000 | 1.0 – 1.1 |
| **1d** | polymer beads, size sort | 30 | x | x | glass | 40x | 220 – 300 | 0 – 0.01 | x | x | 1.0 – 1.1 |
| **1d** | PAA beads, def sort | 20 | x | x | $LiNbO_3$ | 20x | 95 – 105 | 0 – 0.016 | x | x | 1.0 – 1.1 |
| **2d** | blood, RBC sort | 20 | x | x | glass | 40x | 25 – 65 | 0.16 – 0.40 | x | x | 1.0 – 1.1 |
| **2e** | blood, ly sort | 20 | x | x | glass | 40x | 25 – 45 | 0 – 0.10 | x | x | 1.0 – 1.1 |
| **2f** | blood, my sort | 20 | x | x | glass | 40x | 53 – 120 | 0 – 0.15 | x | x | 1.0 – 1.1 |
| **3** | blood, neu sort | 20 | x | FL-2: 10%; FL-3: 20% | glass | 40x | 56 – 100 | x | 75 – 78 | x | 1.0 – 1.1 |
| **4** | blood, neu sort AI | 20 | x | FL-2: 10%; FL-3: 20% | glass | 40x | x | x | x | x | 1.0 – 1.2 |
| **Supplementary Figures** | | | | | | | | | | | |
| **3a** | PAA beads; Def/FL-1 sort | 20 | FL-1: 30% | FL-1: 30% | $LiNbO_3$ | 20x | x | 0.01 – 0.02 | x | 1000 – 10000 | 1.0 – 1.1 |
| **3b** | Polymer beads, br/size sort | 30 | x | x | glass | 20x | 160 – 240 | x | 75 – 85 | x | 1.0 – 1.1 |
| **4** | HL60/S4, Kc167, size sort | 20 | x | x | glass | 20x | 25 – 77 | 0 – 0.15 | x | x | 1.0 – 1.1 |
| **5b** | RBC, high def sort | 20 | x | x | $LiNbO_3$ | 20x | x | 0.15 – 0.40 | x | x | 1.0 – 1.2 |
| **5c** | RBC, low def sort | 20 | x | x | $LiNbO_3$ | 20x | x | 0 – 0.10 | x | x | 1.0 – 1.2 |

**Supplementary Table 4| Sizes of ROIs used for image acquisition.** Depending on the objective used and the width of the measurement channel, the exact ROI size was adjusted to the values listed below. In the case of 20× objective, pixel size corresponds to 0.68 µm and for 40× objective to 0.34 µm.

| | **20× objective** | **40× objective** |
|---|---|---|
| **20-µm channel** | 100 × 40 pixels<br>(68 × 27.2 µm) | 250 × 80 pixels<br>(85 × 27.2 µm) |
| **30-µm channel** | 100 × 45 pixels<br>(68 × 30.6 µm) | 250 × 100 pixels<br>(85 × 34 µm) |

**Supplementary Table 5| Details regarding post-processing of data.** For plotting, the datasets were filtered for area ratio and size. In selected scatter plots, down-sampling to a specific number of points (downs), in case it was exceeded in the dataset, was performed to facilitate distinguishing of different populations. For calculation of purity, only a size filter for small objects was applied, allowing to exclude cell debris. The number of cells used for the calculations of purity in initial and target samples is also indicated. PAA – polyacrylamide, def – deformation, br – brightness.

| | | filters for plotting | | | purity calculation | | |
|---|---|---|---|---|---|---|---|
| | **sample** | **size ($\mu m^2$)** | **area ratio** | **downs** | **size filter ($\mu m^2$)** | **# cells initial** | **# cells target** |
| **Figures** | | | | | | | |
| **1d** | PAA beads, FL-1 sort | 50 – 500 | 1.0 – 1.1 | 6000 | > 25 | 50,138 | 19,195 |
| **1d** | polymer beads, size sort | 50 – 500 | 1.0 – 1.1 | 6000 | > 25 | 19,842 | 6,644 |
| **1d** | PAA beads, def sort | 50 – 500 | 1.0 – 1.1 | 6000 | > 25 | 64,409 | 2,986 |
| **2d** | blood, RBC sort | 23 – 120 | 1.0 – 1.1 | none | > 23 | 45,283 | 12,101 |
| **2e** | blood, ly sort | 23 – 120 | 1.0 – 1.1 | none | > 23 | 45,283 | 11,749 |
| **2f** | blood, my sort | 23 – 120 | 1.0 – 1.1 | none | > 23 | 45,283 | 19,013 |
| **3** | blood, neu sort | 23 – 120 | 1.0 – 1.1 | none | > 23 | 18,220 | 4,185 |
| **4** | blood, neu sort AI | 23 – 120 | 1.0 – 1.2 | none | > 23 | 10,113 | 9,438 |
| **Supplementary Figures** | | | | | | | |
| **3a** | PAA beads; def/FL-1 sort | 50 – 200 | 1.0 – 1.3 | 6000 | > 25 | 48,779 | 1,559 |
| **3b** | Polymer beads, br/ size sort | 50 – 200 | 1.0 – 1.1 | none | > 25 | 6,816 | 3,478 |
| **4** | HL60/S4, Kcl167, size sort | 50 – 200 | 1.0 – 1.1 | 20,000 | > 20 | 62,386 | 3,753 |
| **5b** | RBC, high def sort | 23 – 50 | 1.0 – 1.2 | 20,000 | > 23 | 28,871 | 9,981 |
| **5c** | RBC, low def sort | 23 – 50 | 1.0 – 1.2 | 20,000 | > 23 | 28,871 | 11,177 |

**Supplementary Table 6| Numbers of different blood cell type images used for the selection of the most robust DNN architecture.**

| Class | Training | Validation |
|---|---|---|
| **debris** | 69,959 | 770 |
| **lymphocytes** | 363,936 | 492 |
| **RBCs** | 207,071 | 950 |
| **doublets** | 93,031 | 1,063 |
| **eosinophils** | 110,201 | 1,094 |
| **monocytes** | 211,808 | 1,074 |
| **neutrophils** | 2,349,493 | 3,388 |

**Supplementary Table 7| Numbers of blood cell images from different classes used for DNN training and validation prior to a sorting experiment.**

| Class | Training | Validation |
|---|---|---|
| **CD66+** | 9,030 | 1,951 |
| **CD14+** | 1,451 | 331 |
| **CD14−/CD66−** | 23,493 | 3,555 |
| **CD14+/CD66+** | 232 | 88 |

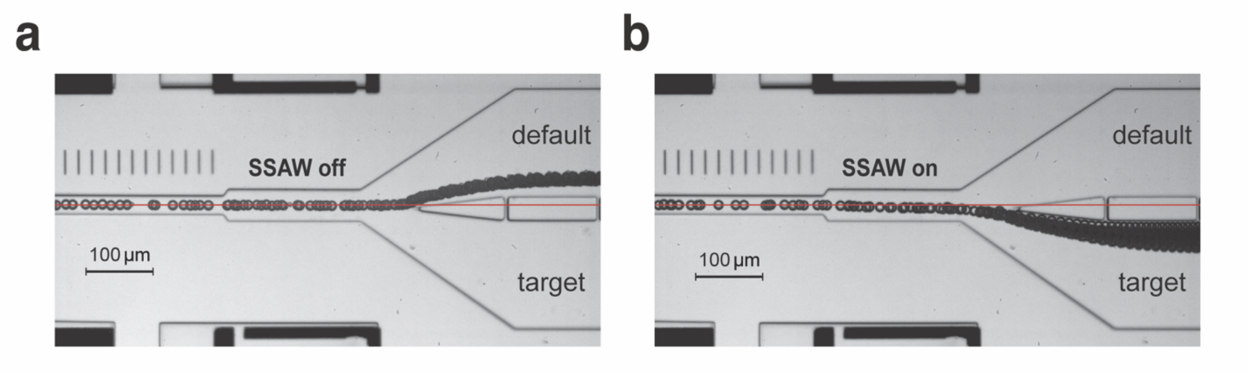


**Supplementary Figure 1| Visualization of bead trajectories in the soRT-FDC chip in the presence and absence of SSAW actuation.** (**a–b**) An image of soRT-FDC chip showing trajectory taken by beads when SSAW actuation is off (**a**) and when SSAW actuation is on (**b**). The images were created by a minimum intensity projection of 700 frames taken over 350 ms. The flow direction is from left to right. The red line indicates the center of the channel. Note the slight off-center position of the bifurcation point.

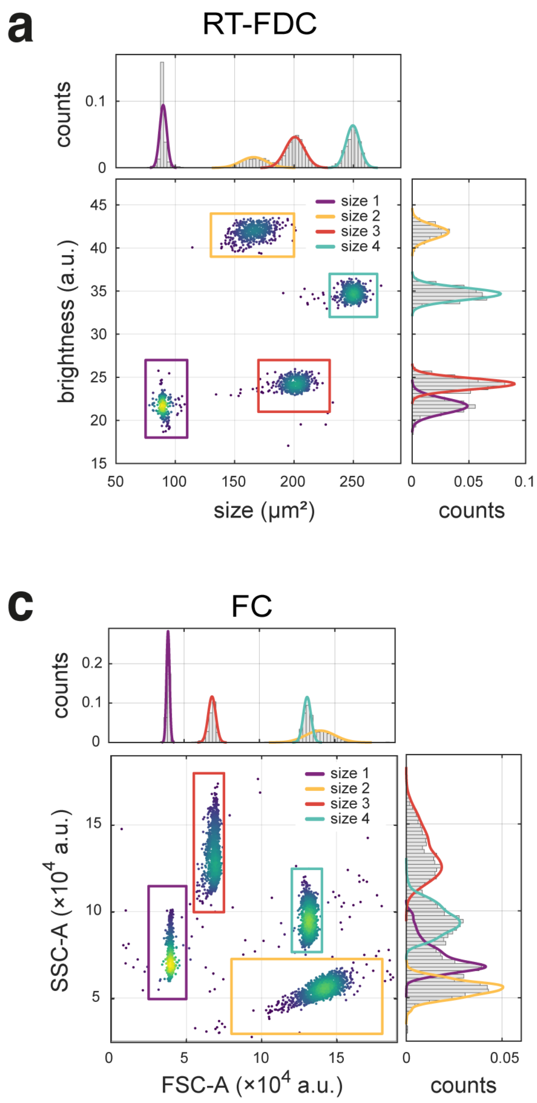


**d**

| | size 1 | size 2 | size 3 | size 4 |
|---|---|---|---|---|
| material | melamine | glass | polystyrene | PMMA |
| | diameter (µm), $\mu \pm \sigma$ | | | |
| supplier specification | 9.78±0.15 | 13.79±0.59 | 15.21±0.31 | 17.23±0.24 |
| | diameter (µm), recalcuated from measured area, $\mu \pm \sigma$ | | | |
| RT-FDC | 10.68±0.17 | 14.53±0.47 | 16.01±0.31 | 17.82±0.21 |
| soRT-FDC | 10.60±0.29 | 14.54±0.54 | 15.71±0.32 | 17.63±0.26 |
| | FSC-A × 10⁴ (a.u.), $\mu \pm \sigma$ | | | |
| FC | 3.93±0.12 | 14.07±1.02 | 6.85±0.25 | 13.16±0.25 |

**Supplementary Figure 2| Determination of bead sizes using RT-FDC, soRT-FDC, and flow cytometery (FC).** A mixture of beads of four different sizes, each made of different material, was analyzed with (so)RT-FDC and FC (BD LSR II, BD Biosciences). We gated for the different bead populations using two-dimensional analysis combining size and brightness in RT-FDC (**a**) and soRT-FDC (**b**), and forward scatter area (FSC-A) and side scatter area (SSC-A) in FC (**c**). In **a**–**c** color map in scatter plots represents event density, the histograms of area and FSC-A are shown on top of scatter plots, and the histograms of brightness and SSC-A on the right-hand side of the scatter plots. Color-coded solid lines represent Gaussian fits to the gated populations of beads. For illustration purposes, data in **a**–**c** was resampled to equalize the relative content of each bead population. (**d**) A summary of bead properties, including bead material, and bead sizes as specified by the supplier, measured with RT-FDC (using 40 × objective and glass substrate) and soRT-FDC (using 20× objective, lithium niobate substrate and an extra polarizer), as well as measured with FACS. Note that the bead size determined by FC is a dimensionless value and not a physical dimension. Furthermore, the relative bead sizes determined with FC do not follow the size order specified by the supplier, as indicated by increasing grey levels in the background.

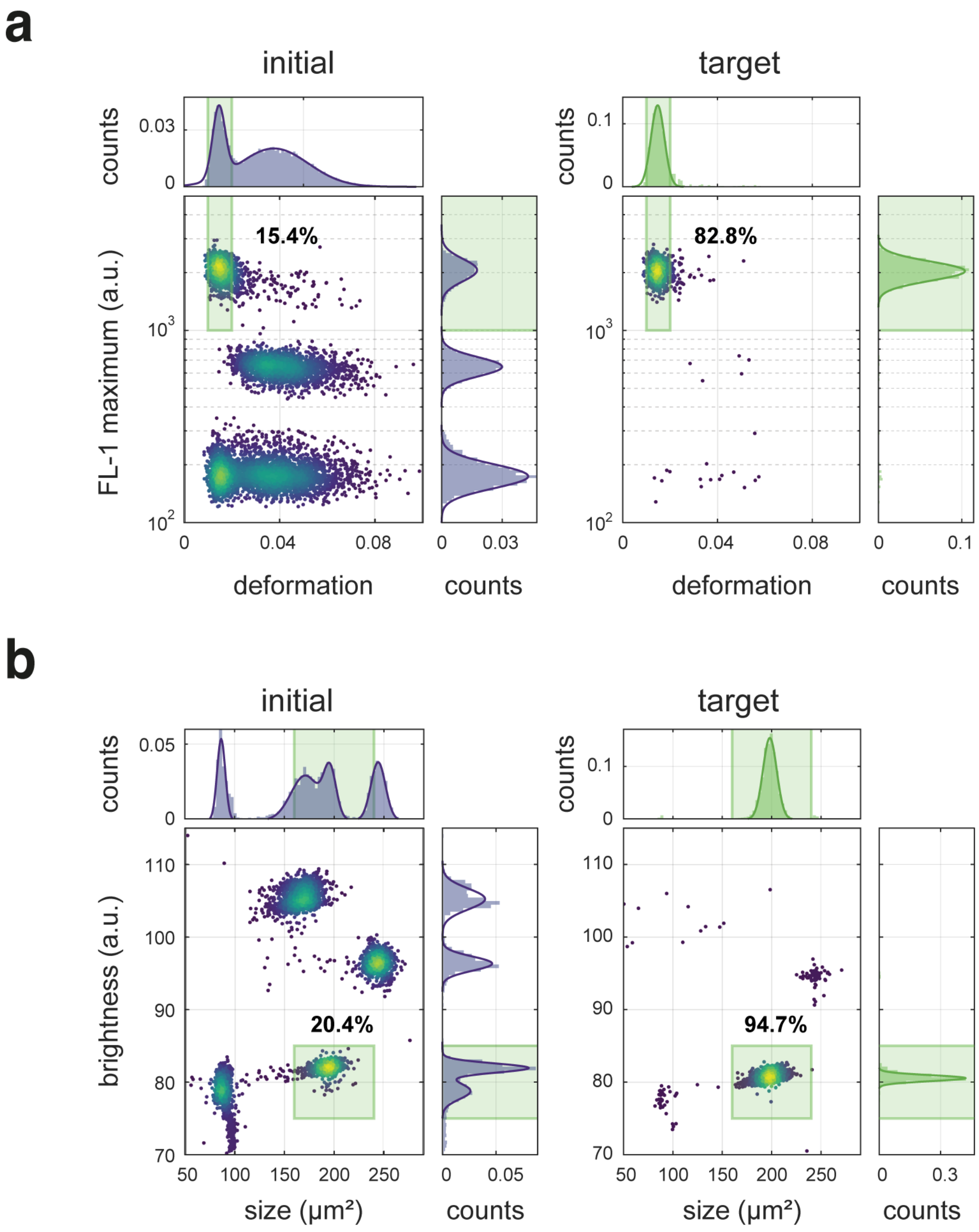


**Supplementary Figure 3| Further examples of bead sorting with two-dimensional gates performed using soRT-FDC.** (**a**) Sorting for deformation together with maximum fluorescence intensity in a mixture of fluorescent and non-fluorescent polyacrylamide beads with different mechanical properties. (**b**) Sorting for brightness and size in a mixture of beads with different diameters, each made of different material (see **Supplementary Figure 2d**). The color map in scatter plots represents event density. The histograms of features presented in the scatter plots are shown on top and on the right of the corresponding scatter plots, the histograms were fit with superpositions of Gaussian functions (solid lines). The gates used for sorting are outlined in green. Percentages on scatter plots indicate the fraction of beads in the sorting gate.

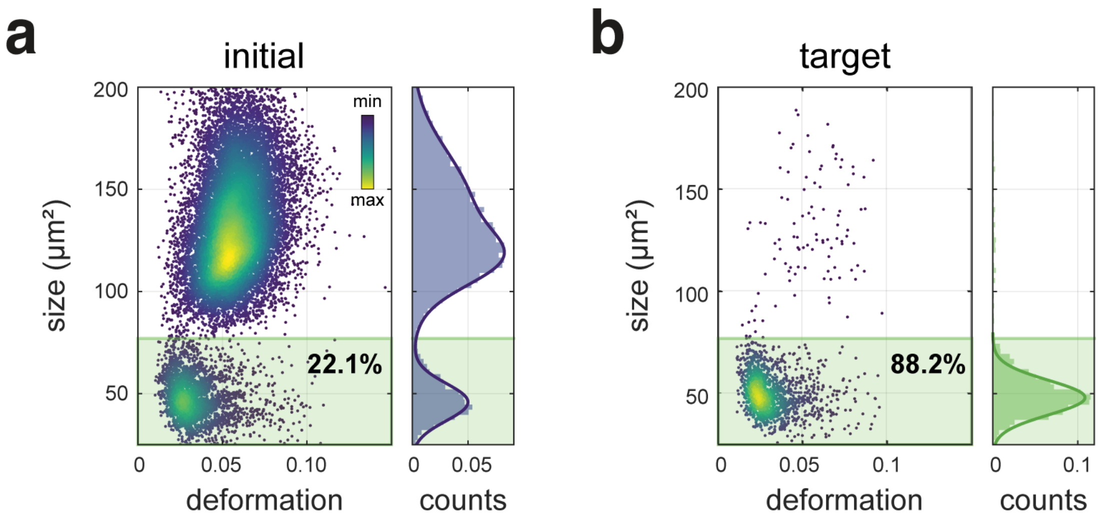


**Supplementary Figure 4| Size-based sorting of a mixture of two cell lines.** (**a**) Deformation-size scatter plot of a 1:4 mixture of Kc167 (smaller size) and HL60/S4 (bigger size) cells in the initial mixture. (**b**) Deformation-size scatter plot of sample collected in the target. Kc167 cells were enriched 4-fold to 88.2%. The green shaded rectangle in both plots indicates the sorting gate (size 25 – 77 µm$^2$, deformation 0 – 0.15). The color map in scatter plots represents event density. The histograms of features presented in the scatter plots are shown on top and on the right of the corresponding scatter plots, the histograms were fit with superpositions of Gaussian functions (solid lines). The gates used for sorting are outlined in green. Percentages on scatter plots indicate the fraction of beads in the sorting gate.

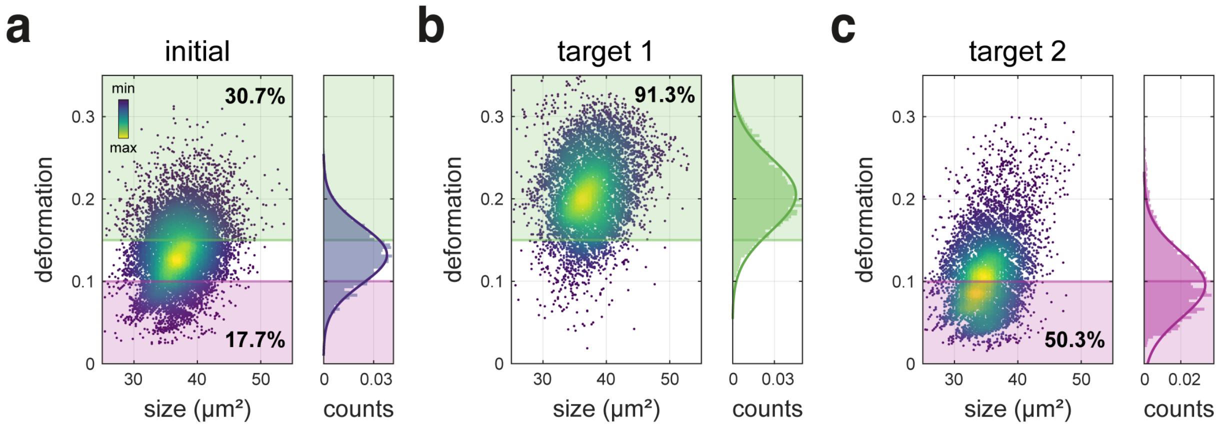


**Supplementary Figure 5| Deformation-based sorting of RBCs.** (**a**) Deformation-size scatter plot for initial measurement of whole diluted blood. The colored rectangles indicate the gates used for RBCs with high (green gate) and low (magenta gate) deformation during two subsequent sorting experiments. (**b**–**c**) Deformation-size scatter plot of sample collected in the target when sorting for cells with low deformation (**b**), 3.0-fold enrichment and 91.3% purity was observed in this case, and when sorting for cells with low deformation (**c**), 2.8-fold enrichment and 50.3% purity was obtained. The color map in scatter plots represents event density. The histograms of features presented in the scatter plots are shown on top and on the right of the corresponding scatter plots, the histograms were fit with superpositions of Gaussian functions (solid lines). The gates used for sorting are outlined in green and magenta, respectively. Percentages on scatter plots indicate the fraction of cells in the sorting gate.

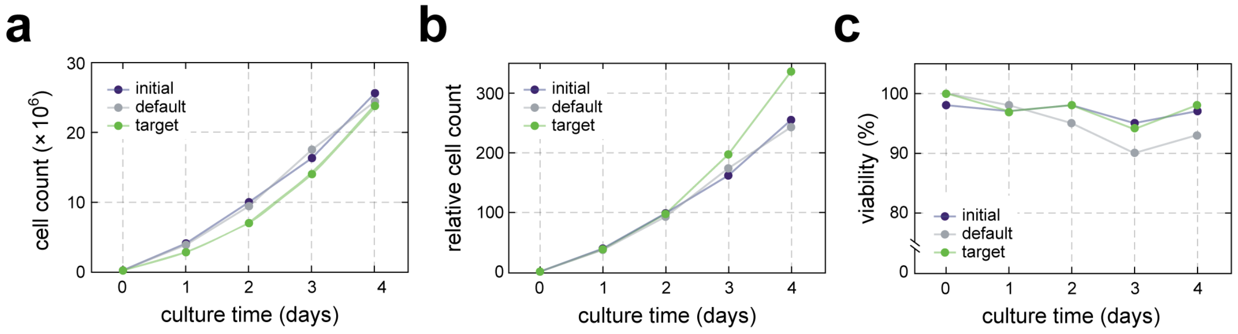


**Supplementary Figure 6| Proliferation and viability of HL60/S4 cells after exposure to SSAW in a soRT-FDC experiment.** (**a**–**b**) Count of cells prepared as for soRT-FDC experiment but not loaded onto the chip (initial), cells run through soRT-FDC chip but not exposed to SSAW (default), as well as cells exposed to SSAW that were collected in the target outlet (target) cultured over 4 days expressed as absolute cell count (a), and relative count with respect to the number of seeded cells (**b**). The number of seeded cells was equal to 70,400 for target, and 100,000 for both initial and default samples. (**c**) Viability of cells in cultures described in (**a**–**b**) evaluated by trypan blue staining.

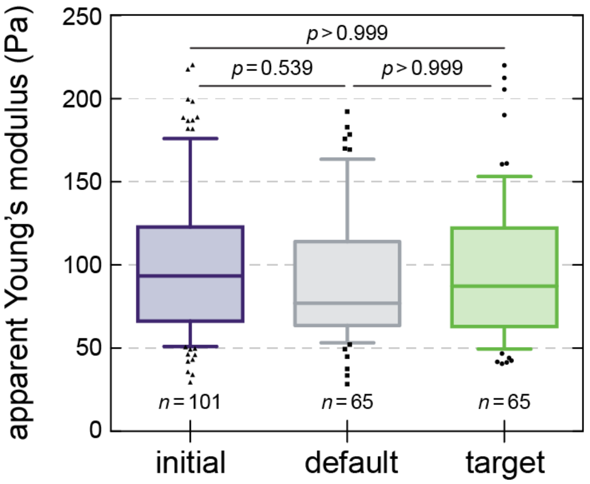


**Supplementary Figure 7| AFM indentation measurement of HL60/S4 cell stiffness after exposure to SSAW in a soRT-FDC experiment.** Box plots of the apparent Young's modulus measured for initial, default, and target samples by AFM. The initial sample was prepared as for the soRT-FDC experiment but not loaded onto the chip, the default sample was collected in the default outlet, i.e., run through the sorting chip but not exposed to SSAW, and the target sample corresponds to cells exposed to SSAW. The number of cells analyzed for each condition, *n*, is indicated in the plot. Boxes extend from 25th to 75th percentiles, with a line at the median. Whiskers span $10^{th}$ to $90^{th}$ percentile. Scattered data points correspond to outliers. Datasets were compared using a Kruskal-Wallis test with a Dunn's multiple comparisons test. The obtained *p*-values are reported in the figure.

**Supplementary Video 1| Size-based sorting of a mixture of Kc167 and HL60/S4 cell lines using soRT-FDC.** The video presents a screen capture of the sorting software window taken during the experiment. The gate of cross-sectional area 25 – 77 μm$^2$ was set to select for the smaller cells (Kc167) (see **Supplementary Figure 4**). The top panel presents an image of the sorting chip. The scale marks in the lower right are spaced 20 μm apart. The analysis ROI is marked with a white rectangle. The SSAW sorting region as well as the target and default outlets are indicated with text labels. Before the sorting start at around 4 s, all cells are directed into the default outlet. Upon sorting start, some of the cells are deflected towards the target outlet. Below the chip image, the bright field images of cells taken in the ROI are shown only for cells classified for sorting. Next to it, the thresholded and binarized images of all detected cells are displayed. The control panel on the lower left shows several real-time metrics, including total number of sorted cells ('Cells sorted'), total number of detected cells ('Cells detected'), the number of detected cells per seconds ('Cells detected [1/s]'), and the number of sorted cells per second ('Cells sorted [1/s]'). The real-time scatter plot of cell size (area) vs deformation of all detected cells is shown on the right-hand side and restarted after ca 500 events. This screen capture was recorded with VLC media player at approximately 25 frames per second, 100 times lower than the frame rate used for analyzing the cells.

**Supplementary Video 2| Brightness-based sorting of neutrophils from RBC-depleted blood using soRT-FDC.** The video presents a screen capture of the sorting software window taken during the experiment. The gate of brightness (75 – 78) and cross-sectional area (56 – 100 μm$^2$) were set to select for the neutrophils (see **Figure 3**). The top panel presents an image of the sorting chip. The scale marks in the lower right are spaced 20 μm apart. The analysis ROI is marked with a white rectangle. The SSAW sorting region as well as the target and default outlets are indicated with text labels. Before the sorting start at around 4 s, all cells are directed into the default outlet. Upon sorting start, some of the cells are deflected towards the target outlet. Below the chip image, the bright field images of cells taken in the ROI are shown only for cells classified for sorting. Next to it, the thresholded and binarized images of all detected cells are displayed. The control panel on the lower left shows several real-time metrics, including total number of sorted cells ('Cells sorted'), total number of detected cells ('Cells detected'), the number of detected cells per seconds ('Cells detected [1/s]'), and the number of sorted cells per second ('Cells sorted [1/s]'). The real-time scatter plot of brightness vs deformation of all detected cells is shown on the right-hand side and restarted after ca 500 events. This screen capture was recorded with VLC media player at approximately 25 frames per second, 100 times lower than the frame rate used for analyzing the cells.
